# Roseobacter enrichment early in life facilitates future colonization of Roseobacter bacteria and improves long-term survival against *Vibrio aestuarianus* in the Pacific oyster

**DOI:** 10.64898/2026.01.09.698610

**Authors:** Marissa D. Wright-LaGreca, Andrew H. Loudon, Amanda E. Bates, Denman M. S. Moody, James A. Dennis-Orr, Korrina M. Gilchrist, Fletcher E. Falk, Timothy J. Green

## Abstract

Exposure to beneficial bacteria during early immune development may promote long-term survival against pathogens, known as “microbial education.” As marine diseases intensify with ocean warming, methods to improve marine organism’s robustness against disease will be valuable for mitigation. Here, we experimentally tested whether exposure to high temperature seawater, seawater enrichment with Roseobacter bacteria, or a combination of both during the first 24 hours of life enhances long-term survival of the Pacific oyster (*Crassostrea gigas*) against the widespread marine pathogen, *Vibrio aestuarianus* subsp. *francensis,* at high temperatures (24 °C). Exposure to high temperatures early in life did not improve future survival during disease challenges and caused high larval mortality. Conversely, Roseobacter enrichment during high temperature exposure resulted in a “microbial rescue effect,” improving larval survival. We found that Roseobacter enrichment during the first 24 hours of life improved future survival against *V. aestuarianus* at high temperatures by up to ∼28% at 13 days post-fertilization (dpf) and ∼30% at 90 dpf. Ultimately, the supplemented Roseobacter species did not remain associated with the host microbiome but instead was replaced by a high abundance of other Roseobacter bacteria at 90 dpf. These findings suggest that early Roseobacter enrichment facilitates future colonization by other Roseobacter species, which may in turn protect against pathogenic *V. aestuarianus*, supporting the concept of microbial education in marine invertebrates and its potential use to combat marine diseases.

**Importance:** This research identifies key aspects of early developmental processes in a marine invertebrate, *Crassostrea gigas*, and explores how these processes can be utilized to alter host-microbe interactions later in life.

## 2. Introduction

Host-associated microbes can be vital to host health, including defence against pathogens (McFall-Ngai et al., 2013). During early animal development, immune processes are activated and shaped by surrounding microbiota (McFall-Ngai et al., 2013; Gensollen et al., 2016). Exposure to microbes early in life trains the immune system, a phenomenon known as “microbial education”, improving future survival against disease (Datan et al., 2024). It is hypothesized that microbial education enhances immune function by promoting the host’s recognition and response to pathogens, and by teaching the host to sequester beneficial microbes that compete against pathogens (Nyholm and Graf, 2012; Desriac et al., 2014; Datan et al., 2024). The absence of “teaching” microbes early in life can increase susceptibility to disease in animals (Rook et al., 2005; Sorée et al., 2022; Fallet et al., 2022). Rising ocean temperatures enhance pathogen impacts through temperature-dependent mechanisms, including the activation of virulence genes, increased proliferation, and spread to new geographic regions (Harvell et al., 1999; Burge & Hershberger, 2020). Furthermore, extreme temperatures can cause physiological stress to the host, thereby reducing its immune capacity and increasing disease susceptibility (Lannig et al., 2006). As a result, rising ocean temperatures can increase the frequency and intensity of marine diseases while also making marine organisms less adept at defending against them. Microbes in the host’s microbiome can serve as a lens through which ecological conditions, including abiotic (e.g., high temperatures) and biotic factors (e.g., pathogens), affect host health (Kohl, 2025). Host-associated microbes can cause a ‘microbial rescue effect’ by stabilizing host health and preventing mortality (Lehtinen, 2023). The addition of bacteria that benefit host health during the microbial education period may enhance microbiome composition and improve survival against pathogens in marine organisms, providing a tool to protect marine organisms amid rising ocean temperatures.

Recently, the Pacific oyster (*Crassostrea gigas*, also known as *Magallana gigas*) has emerged as a model species for studying microbial education in marine invertebrates (Fallet et al., 2022; Mackenzie et al., 2022; Sorée et al., 2022; Datan et al., 2024). This is in part due to the controlled conditions in which *C. gigas* can be propagated (in aquaculture hatcheries), where the microbial environment can be manipulated before transfer to open-ocean conditions during the juvenile life stages, exposing them to dynamic environmental conditions and pathogens. Invertebrates such as *C. gigas* lack an adaptive immune system capable of developing antibodies against specific microbes and instead rely on an innate immune system (i.e., non-specific, first-line defences); however, our understanding of the innate immune system’s capacity for adaptive-like immune memory responses is growing (Bekkering et al., 2021). For example, *C. gigas* has a highly polymorphic genome with high levels of post-transcriptional/translational modifications, which enables a diverse array of receptors to detect specific bacteria (Huang et al., 2015). As filter feeders with an open circulatory system, *C. gigas* are in direct contact with the external microbiota, yet their microbiome composition can differ markedly from that of their surroundings, suggesting selection over microbial colonization (Unzueta-Martínez et al., 2022). Collectively, these features make *C. gigas* an ideal model for exploring microbial education processes and their role in long-term survival against disease in marine systems.

Over recent decades, mass mortalities of *C. gigas* have been linked to extreme temperature events and subsequent infection by pathogenic *Vibrio* bacteria (Garnier et al., 2007; Green et al., 2019; Cowan et al., 2024). The oyster microbiome typically contains bacteria within the genus, *Vibrio*; however, some are opportunistic pathogens, where a clear onset of pathogenicity ensues during environmental stressors, such as high temperatures (de Lorgeril et al., 2018). *Vibrio aestuarianus* is a well-known pathogen of juvenile (Cousson et al., 2025) and adult oysters (Garnier et al., 2007) and has been a growing concern in the North Pacific area (Cowan et al., 2023 and 2024). Previous research has shown correlations between *V. aestuarianus* abundance and increasing sea surface temperatures (Cowan et al., 2023 and 2024). In addition to external *Vibrio* bacteria becoming more numerous in the environment, empirical evidence suggests that *Vibrio* occupying the microbiome of healthy oysters proliferate and become virulent within the host at high temperatures (Green et al., 2019).

Members of the Roseobacter bacterial group can inhibit the growth of *Vibrio* pathogens (Brinkhoff et al., 2004; Bruhn et al., 2005). *Roseobacter* is a genus containing only a few species (e.g., *Roseobacter litoralis* and *Roseobacter denitrificans*), but, more broadly, Roseobacter is used to classify a group of ∼68 bacterial genera and 164 species with distinct characteristics and/or phylogenetic relatedness (Pujalte et al., 2014; Simon et al., 2017; Liang et al., 2021). Several Roseobacter bacteria have symbiotic relationships with phytoplankton such as diatoms, dating back 260 MYA (Luo et al., 2014). Phytoplankton supply nutrients such as dimethylsulfoniopropionate (DMSP), and in exchange, Roseobacter provide vitamins and protection from pathogens through the production of potent antibacterials such as tropodithietic acid (TDA) (Liang, 2003; Luo et al., 2014). Filter feeders that feed on phytoplankton, such as *C. gigas*, are likely to be in close contact with phytoplankton-associated Roseobacter bacteria and may have a long-standing evolutionary relationship with these bacteria. Roseobacter are associated with the microbiome of bivalves throughout life-history stages (i.e., larval to adult) and may represent important and persistent microbes within the *C. gigas* microbiome (Pierce and Ward, 2019).

Past studies have found a protective effect of Roseobacter bacteria (*Phaeobacter gallaeciensis*) for host survival against *Vibrio* pathogens (e.g., scallop Ruiz-Ponte et al., 1999; e.g., shrimp Balcázar et al., 2007); however, these beneficial effects only last for hours after supplementation (e.g., oyster Karim et al., 2013; e.g., oyster and scallop Kesarcodi-Watson et al., 2012), suggesting the supplemented bacteria are not incorporated into the host’s microbiome. Immune capabilities are present as early as 4 hours post-fertilization (blastula stage) (Song et al., 2016), and thus, it may be best to supplement bacteria as early as possible (e.g., during fertilization), when major immune processes are still forming; however, some research has found that high bacterial loads during the first hours of life results in moderate to severe developmental abnormalities (Douillet & Langdon, 1993; Datan et al., 2024). Therefore, an important consideration when testing the effects of microbial education is whether early survival is impacted. Overall, our understanding of microbial education and its use as a tool for disease prevention in marine organisms is nascent and requires much more research.

Here, we tested the microbial education concept by exposing the model host, *C. gigas*, to potentially protective bacteria (Roseobacter) early in development and subsequently assessed whether host survival against a pathogen (*Vibrio aestuarianus*) and microbiome composition were altered later in life. Additionally, we tested whether early-life exposure to an abiotic stressor (high temperatures) further enhances long-term survival against disease during high temperatures. Specifically, we exposed *C. gigas* to Roseobacter-enriched (*Celeribacter baekdonensis*) and/or high temperature seawater during the first 24 hours of life and subsequently, their survival against *Vibrio aestuarianus* subsp. *francensis* and high temperatures were assessed. We predict that early Roseobacter enrichment and exposure to high temperatures will improve future survival during *V. aestuarianus* and high temperature challenges. Additionally, we predict that changes in survival may co-occur with long-term microbiome changes, including increases in the abundance of beneficial bacteria (i.e., symbionts) that may protect against pathogens, supporting the microbial education hypothesis. This research sheds light on the intersection between early environment, immune development, and microbiome formation, and its potential effects on host-pathogen and host-symbiont interactions.

## 3. Methods

### 3.1 Experimental design

To test the effects of Roseobacter exposure and high temperatures we used a factorial design, where larvae were exposed to either control (18.2 °C ± 0.321), Roseobacter-enriched (18.4 °C ± 0.50), High temperature (HT) (25.7 °C ± 0.884), or Roseobacter-enriched + HT (27.9 °C ± 2.11) seawater for the first 24 hours of life. Treatments were replicated across the four genetic lines (denoted pair-mated “families”) of *C. gigas*. Replicating across different oyster genetic lines was chosen because larval survival and microbiome differed between genetic lines in pilot studies, and this has been noted in other published studies (King et al., 2019). Four genetic lines ensured at least three family replicates were available if one was not compatible (i.e., low fertilization or survival). Since dams (mothers) are the driving force in larval survival and microbiome composition (Unzueta-Martínez et al., 2022), we chose four dams to be fertilized with a single sire (father). We recognize the limitations of having a true sample size of one tank (n = 1) for each treatment and family cross, but we were also cautious of using one family across multiple treatment tanks. Such experimental designs can be problematic if survival and microbiome composition is highly variable between families, causing a “family bias.” Ideally, the four families would be replicated in triplicate tanks across all experimental treatments (a total of 48 tanks); however, research hatcheries are limited by time, space, and equipment required for such experiments.

For the first 24 hours, fertilized eggs were placed in 20 L buckets with seawater, aerating tubes, and aquarium heaters. After 24 hrs, the larvae were obtained using 48 µm mesh sieves and placed in 250 L rearing tanks for the remainder of development. Bacterial water and materials were then disinfected using a 1% v/v bleach solution. Sampling points for larval enumeration and microbiome analysis (blue asterisks) and for *Vibrio aestuarianus* + high temperature disease challenges (red asterisks) across life-stages are shown in Figure 1. Treatment exposure and larval rearing took place at the Vancouver Island University (VIU) Deep Bay Marine Field Station (49.457N, -124.733W) research hatchery.

**Figure 1.**
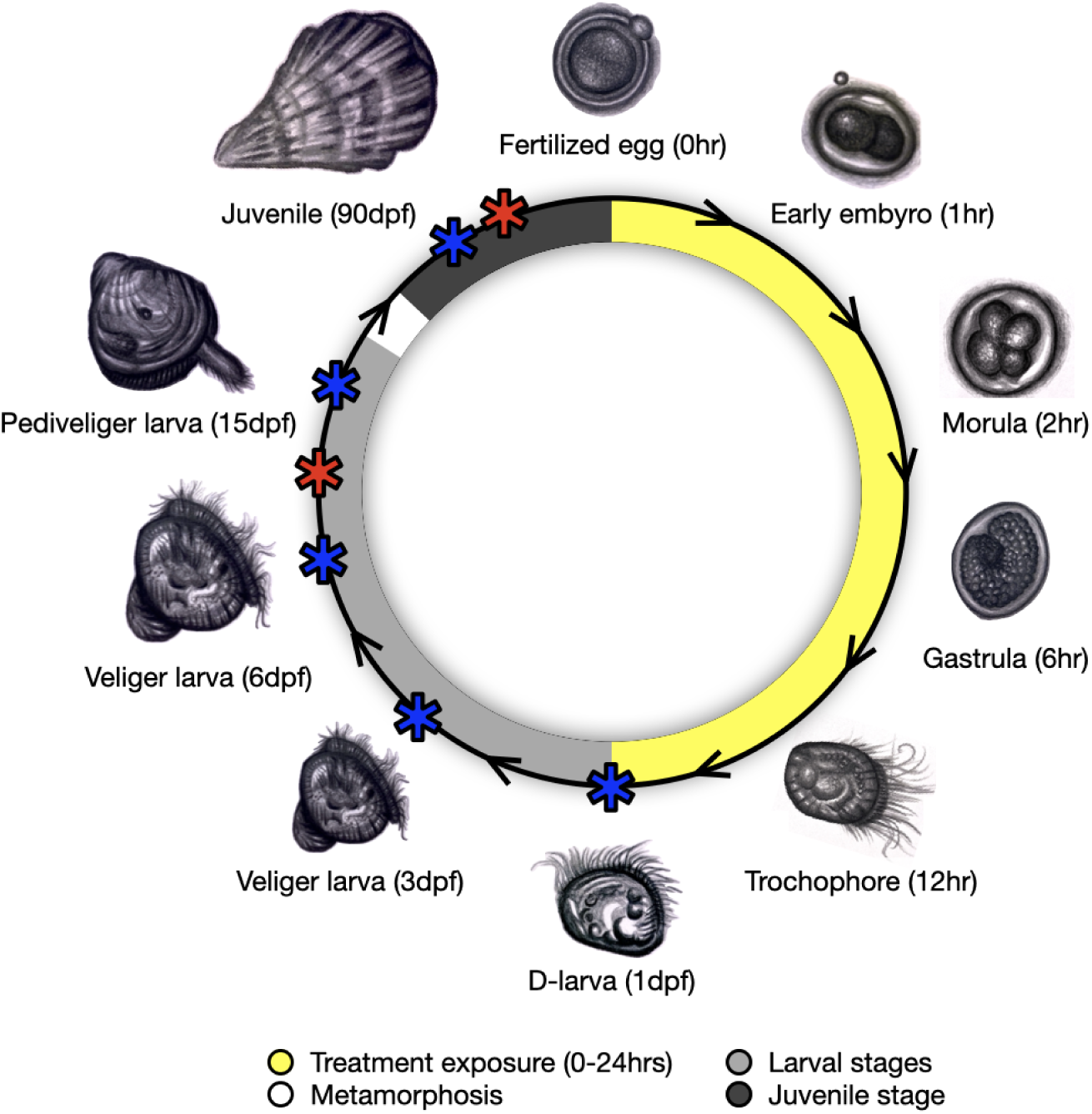
Illustration of oyster development during treatment exposure (first 24 hours of life) and of sampling points in subsequent life-stages (larval and juvenile) for larval enumeration and/or microbiome analysis [blue asterisks; 1, 3, 6, 15, and 90 dpf (days post-fertilization)] and *Vibrio aestuarianus* + high temperature disease challenges (red asterisks; 13 and 90 dpf). hpf = hours post-fertilization. Larval images illustrated by Bri Cooper.

### 3.2 Roseobacter isolation & whole-genome sequencing

The Roseobacter bacterium was isolated from healthy *C. gigas* larval samples at the VIU Deep Bay Marine Field Station in the summer of 2018 by streak isolation on Marine Agar (Sigma-Aldrich, Millipore #76448) culture plates. To determine the identity of the bacteria, the DNeasy Blood and Tissue kit (QIAGEN: #69504) was used for DNA extraction. Whole genome sequencing (WGS) using a shotgun sequencing method for probiotic identification was conducted on the Illumina NovaSeq platform [150 base pair (bp) paired-end reads with 60 million sequence depth] at the Genome Quebec Sequencing Centre (Montreal, Canada). Read quality assessment was performed with FastQC (v12.1) (Andrews, 2010). Low-quality reads (Phred score < 30) and adaptors were removed using Trimmomatic (v3.9) (Bolger et al., 2014). Genome assembly was conducted with Unicycler v0.5.1 (optimal kmer = 127) (Wick et al., 2017). To identify the isolate at a species level, the 16S rRNA gene was extracted from the WGS and aligned against the National Center for Biotechnology Information (NCBI) nucleotide Basic Localized Alignment Search Tool (nBLAST), matching with *Celeribacter baekdonensis* (top accession #: #CP028475). Following this, the probiotic genome assembly was aligned against a complete chromosome assembly of *C. baekdonensis* using the FastANI program (v1.34) (Jain et al., 2018). Genome alignments were repeated using closely related reference genomes (*Celeribacter neptunius*, *Celeribacter marinus*, *Alliroseovarius crassostrea*, *Roseobacter ponti*, *Roseobacter fuci*) and a distantly related reference genome (*Escherichia coli*) for comparison. Alignment matches of ≥ 95% were used as a species-match threshold (Rodriquez-R et al., 2024).

### 3.3 Roseobacter growth & preparation

*Celeribacter baekdonensis* was grown under two conditions: shaking at 50 rotations per minute (RPM) and non-shaking, both at 20 °C for four days in 500 mL Erlenmeyer flasks containing 200 mL of marine broth (Sigma-Aldrich, Millipore #76448). The shaking and non-shaking conditions were implemented as previous studies have suggested a potential effect of shaking conditions on antimicrobial activity (Bruhn et al., 2007). On the day of spawning, *C. baekdonensis* cells were concentrated by centrifugation and washed with sterile seawater to remove nutrient broth. Cultures from both culture conditions (shaking and non-shaking) were pooled into a single 50 mL falcon tube to achieve a final concentration (OD_600_) of 2.10. Probiotic treatments received 4 mL of the prepared culture that was diluted in the 20 L buckets filled with 5 μm filtered seawater. To estimate the final concentration of *C. baekdonensis* added, a serial 10-fold dilution series was plated on Marine Agar and colony forming units (CFUs) were counted the following day, indicating a final concentration of 1.8 x 10^2^ cells·mL^-1^ in the 20 L buckets.

### 3.4 Spawning, larval rearing, and sampling

Four *C. gigas* families were made using four females and one male sourced from the VIU oyster breeding program. Broodstock were strip-spawned, and eggs were counted on a Sedgewick Rafter counting chamber and diluted to a concentration of 570 eggs·mL^-1^ to achieve an estimated one million fertilized eggs per tank. Every second day, larvae were caught on 48 μm mesh screens, then tanks were cleaned with a diluted bleach solution (100 ppm) and washed with 5 μm filtered seawater. Before returning larvae to their respective tanks, larvae were sampled for microbiome analysis and to estimate larval survival and growth rate. For microbiome sampling, 10 mL of larval suspension was collected in 15 mL Falcon tubes, centrifuged, and larval pellet was immediately frozen on dry ice and subsequently transferred to a -80 °C freezer until 16S rRNA sequencing preparation. To estimate larval abundance per tank during rearing (reflective of larval survival), on tank cleaning days, triplicate 250 µL larval samples were collected in 0.9 mL Eppendorf tubes, and 100 µL of 10% neutral buffered formalin (NBF) was added to preserve until counting. Additionally, 100 µL of larval solutions were sampled for immediate tank counts and larval size estimates, which were used to determine algal diet requirements (amount and algal species mix). Larvae were fed a mixed diet of microalgae containing *Isochrysis* sp., *Pavlova* sp., and *Chaetoceros calcitrans*, following the feeding recommendations described in “Hatchery culture of bivalves: a practical manual” (Helm et al., 2004).

At 15 days post-fertilization (dpf), larvae had developed eyespots and had high lipid content, and thus, were deemed ready for settlement (metamorphosis). The metamorphosis from free-swimming to non-motile generally marks the transition from the larval to juvenile life-stage (Figure 1). To induce settlement, larvae were incubated in 0.0008 M epinephrine (Sigma-Aldrich: E4375) solution for one hour before being returned to their respective tanks. The following day settled oysters were gently brushed off their tanks and collected on 220 μm mesh screens. Collected oysters were placed in downwellers (i.e., polyvinyl chloride pipe silos with 180 μm mesh screens and seawater flowing from above) in a flow-through nursery system until the *Vibrio aestuarianus* + high temperature challenge at 90 dpf.

### 3.5 Enumeration of tank numbers through larval rearing

To enumerate larvae, 100 µL of vortexed larval-NBF suspension was aliquoted into a 48-well Eppendorf cell-culture plate. The Leica THUNDER Imager (5X magnification) was used to visualize each well with the Spiral Navigation application. An averaged value from four count replicates was used as an estimate of larval tank count for each tank at each time point. A generalized additive model was utilized to account for non-linearity of larval counts through time [effective degrees of freedom (edf) > 1, *p <* 0.01], with the factors of treatment, family, and their interaction. To further investigate differences between treatments, a generalized additive mixed model with oyster family as a random factor was used. All models were fit with a tweedie distribution using the mgcv package in R (Wood, 2011). Model fit was confirmed by visualizing Q-Q and residuals plots using the stats R package (R Core Team, 2025) and using the gam.check function with the mgcv R package (Wood, 2011). All analyses were conducted in R Studio software (version 4.4.3)

### 3.6 Pathogen isolation & disease challenge dosage

The *V. aestuarianus* pathogen was isolated from moribund oysters during an oyster mortality event in 2018 in Baynes Sound, BC (Cowan et al., 2024). Whole genome sequencing was previously performed, and the genome assembly had an ANI match of 99.99% with *Vibrio aestuarianus* subsp. *francensis* (GenBank ID: #GCF_046850195.1), compared to a 97.39% with the other subspecies, *Vibrio aestuarianus* subsp. *aestuarianus* (GenBank ID: #GCF_012395185.1). This isolate has been used as a juvenile oyster pathogen in previous studies (Mackenzie et al., 2022; Surry et al., 2024), and this subspecies is a causative pathogen of *C. gigas* in France (Garnier et al., 2008) and Ireland (Coyle et al., 2023). Furthermore, whole-genome analysis of this isolate confirmed the presence of a complete *varS* gene, in which frameshift mutations eliminate pathogenicity (Goudenège et al., 2015). Although different *Vibrio* pathogens are primarily associated with larval mortality (e.g., *Vibrio coralliilyticus* and *Vibrio tubiashii*; Richards et al., 2015), *Vibrio aestuarianus* subsp. *francensis* was used for consistency to assess disease resilience across multiple life stages (larval and juvenile) and because testing survival against the pathogen associated with summer mortality (i.e., *V. aestuarianus*) was our primary objective.

The *V. aestuarianus* and high temperature laboratory challenges are designed to be short and controlled representations of summer mortality events to assess disease robustness. Empirical evidence suggests infected oysters can shed an average of 7.2 x 10^4^ *V. aestuarianus* cells·mL^-1^ per oyster per day, with some oysters exceeding this (2.0 x 10^6^ cells·mL^-1^·day^-1^), denoted “super-shedders” (Lupo et al., 2019). In field and hatchery settings, oysters are kept in close proximity and thus may be exposed to multiple infected individuals simultaneously. Additionally, planktonic bacterial cells (e.g., *Vibrio* grown in broth) are filtered and ingested at rates up to 50% lower than those of larger, particle-associated cells (e.g., bacteria attached to algal particles), where particle-associated *Vibrio* cells would be more typical in natural environments (Kach & Ward, 2008; Froelich et al., 2013). Thus, the *V. aestuarianus* dose used here (larval disease challenge: 1.72 x 10^8^ cells·mL^-1^ total over 3 days; juvenile disease challenge: 2.80 x 10^8^ cells·mL^-1^ total over 6 days) is not far from potential exposures during the height of a disease event under confined conditions. Dosing *V. aestuarianus* at the upper range of that expected during a disease event avoids holding the oysters in laboratory conditions over an extended time period, as they are expected to die faster at higher concentrations of *V. aestuarianus*. To test if the addition of non-pathogenic bacteria affects the survival of oysters (e.g., by reducing oxygen of the seawater in the wells), a pilot study using these well-assay protocols with *Escherichia coli* was conducted and demonstrated no impact on oyster survival compared to control (no bacteria) conditions (*p =* 0.283).

### 3.7 Larval (13 dpf) *Vibrio aestuarianus* challenge

On 13 dpf (veliger stage), a small subset of larvae was sampled for a *Vibrio aestuarianus* and high temperature challenge (24 °C). Family 4 was not included in any survival analyses [larval and juvenile (section 3.7) *V. aestuarianus* + high temperature challenges] because the control tank for Family 4 died by 13 dpf. The HT treatment was not analyzed in any survival analyses, as only one tank (containing one family) remained. A concentration of 30.3 ± 20.2 larvae·well^-1^ was added to each well. To test if variations in larval concentration per well impacted larval survival, a Pearson correlation test was utilized which indicated no significant relationship between survival and well density (R^2^ = -0.02, *p =* 0.92) (Supplementary Materials, Figure 1). Following the addition of larvae, wells were filled with autoclaved seawater to a total volume of 0.75 mL.

*Vibrio aestuarianus* was grown in Tryptic Soy Broth (Millipore Sigma: #41298) containing 2% NaCl in 50 mL Falcon tubes overnight at 24 °C and 200 RPM. Following this, the bacterial solution was centrifuged, and the cell pellet was washed with 0.2 μm filtered seawater to remove residual broth. The cell pellet was resuspended in seawater to a concentration of OD_600_ = 0.50. To each well, 100 μL of *V. aestuarianus* solution was added to reach a final concentration of OD_600_ = 0.06 (∼1.72 x 10^8^ cells·mL^-1^). The disease challenge lasted 36 hrs, and subsequently, larvae were phenotypically assessed.

Larvae were visualized on the Leica THUNDER Imager (5X magnification) and were characterized as alive (swimming and gut movement visible), alive but non-motile (static and gut movement visible), or dead (static and no gut movement visible). Dead larvae were counted, and subsequently 100 μL of 10% NBF was added to each well to fix larvae for total larval counts. Following total larval counts, the number of live larvae was subtracted to calculate live: dead ratios. Binary survival data were analyzed using a binomial generalized linear model using the stats R package (R Core Team, 2025). Residuals were fitted and visualized with the DHARMa package (Hartig, 2022). Since logistic regression can be sensitive to overdispersion, a Pearson Chi-squared statistic was used to test for this, revealing overdispersion was not present (Chi-squared = 304(296), *p* > 0.05). Following this, we tested if differences in larval survival were explained by treatment, family, or their interaction using an ANOVA (type = II, Survival ∼ Treatment*Family) followed by a pairwise comparison using the Estimated Marginal Means (emmeans) R package (Lenth, 2024) with a Bonferroni *p* adjustment. To further identify if treatment groups had a significant effect on larval survival while accounting for high variability between families, a binomial generalized mixed model (GLMM) with oyster family as a random factor using the glmer function in the lme4 package (Douglas et al., 2015). The GLMM model fit, overdispersion, and post-hoc analysis were tested as described above.

### 3.8 Juvenile (90 dpf) *Vibrio aestuarianus* + high temperature challenge

At 90 dpf (juvenile stage), oysters were placed into 6-well Eppendorf cell culture plates (n = 10 oysters per well) and each well was replicated four times (n = 40 oysters from each tank). *Vibrio aestuarianus* preparation followed the same steps as mentioned above (section 3.5), and the final solution had a concentration of OD_600_ = 0.11 (2.80 x 10^8^ cells·mL^-1^ based on performed *Va* growth curves). Control wells (no bacteria) were filled with 5 mL of 5 μm filtered seawater. One mL of algae was added on the first day of the disease challenge to encourage *C. gigas* filter-feeding and ingestion of *V. aestuarianus* cells. Oysters were incubated at 24 °C. Daily live/dead counts were performed for six days. Mortality was determined by the inability of *C. gigas* to retract their mantle after being prodded with sterile forceps and/or a gaping shell valves, indicating non-active adductor muscles.

The Kaplan-Meier maximum likelihood estimator was used to analyze variations in time of death between treatments with the survival R package (Therneau, 2024) and survival curves were generated with the survminer R package (Kassambara et al., 2024). Estimated hazard ratios relative to the control treatment were generated with the Cox proportional hazards regression model. To account for family-level variation, a mixed-effects Cox model (coxme R package, Therneau, 2024) with family as a random factor was used to test for treatment differences and calculate hazard ratios (95% confidence intervals), which were plotted using the ggplot2 R package. Survival data met the proportional hazard assumption (*p* > 0.05).

### 3.9 16S rRNA library formation & sequencing

Oyster DNA tissue extraction for 16S rRNA sequencing was performed with the QIAGEN DNeasy^®^ Blood & Tissue kit (QIAGEN: # 69504). The 16S Metagenomic Sequencing Library Preparation protocol (Illumina Inc, San Diego, California, USA: #15044223) was largely followed for 16S rRNA sequencing preparation. The PCR protocol was modified to 95 °C for 5 mins, 95 °C for 10 secs, 55 °C for 10 secs, 72 °C for 1 min (steps 2 - 4 repeated 32X), and 72 °C for 5 mins. The first PCR was performed using 5 µL of purified DNA (1:100 dilution) using 16S rRNA (V3/V4 region) primers. PCR products were confirmed on a gel and PCR-cleaned using the QIAquick PCR cleanup kit (QIAGEN: #28104). Indexing PCR was conducted using the iProof High-Fidelity PCR Kit (Bio-Rad Laboratories: #1725330). Again, PCR products were confirmed on a gel and PCR products were purified. Finally, library quantification was performed using the NEBnext^®^ Library Quant kit BioLabs for Illumina® (#E7630S) with a qPCR reaction of 8 µL of the NEB Hot Start 2x Master Mix (#M0496S) and 2 µL of sample diluted 1000-fold. Following library quantification, all samples were diluted to an approximate concentration of 4 ng·µL^-1^ and subsequently pooled into a single 1.5 µL Eppendorf tube. Samples were sent to the University of British Columbia Sequencing & Bioinformatics Consortium for sequencing on the MiSeq sequencing platform.

### 3.10 Microbiome data processing and statistical analyses

The 16S rRNA reads were processed in R Studio (v4.3.3), following the Workflow for Microbiome Data Analysis: from raw reads to community analyses document (Callahan et al., 2016). Taxon were assigned using the Silva classifier v132 at a similarity threshold of 99% and classified as Amplicon Sequence Variants (ASVs) (Yilmaz et al., 2014). Taxa belonging to non-bacterial 16S rRNA sequences (Archaea, Mitochondria, and Chloroplasts) were filtered from the data. Taxa accounting for fewer than 200 total reads were filtered out. Read counts were rarefied to 4660 reads, removing two samples (Sample IDs: T19-1 and T8-6).

To analyze differences in alpha diversity (i.e., number of taxa), Shannon, Chao1, and inverse Simpson diversity indices were compared between treatment, family, and time points, using an Analysis of Variance (ANOVA) test (stats R package, R Core Team, 2025) followed by Tukey’s post-hoc test when parametric assumptions were met, and the Kruskal-Wallis chi-squared test with Dunn’s post-hoc test (dunn.test R package, Dinno, 2024) when data were non-parametric (Alpha diversity ∼ treatment*family*time). To investigate changes in beta diversity across time and between treatment and family groups, a Permutational Multivariate Analysis of Variance (PERMANOVA; Beta diversity ∼ treatment*family) was utilized across each factor combination using the vegan R package (Oksanen et al., 2013). Due to high variation in beta diversity across time points, the factor of time did not pass the homogeneity tests; thus, time points were analyzed separately. To visualize overall changes in beta diversity between treatments across time points, Bray-Curtis similarity matrices followed by Principal Coordinates Analysis (PCoA) with 999 permutations were generated for each time point using the ordinate function in the phyloseq R package (McMurdie and Holmes, 2013). Members of the oyster’s “core” microbiome, here defined as ASVs with a prevalence of 90% or greater across all time points, are hypothesized to play essential roles in the host’s biological functions (Risely, 2020). Thus, core members across time points were analyzed to define persistent ASVs and to detect differences in the core microbiome between treatments.

To determine the length of time that ASVs corresponding to *C. baekdonensis* (ASV7/18) remained in the Roseobacter-enriched treatments, the average relative abundance of *C. baekdonensis* ASVs was plotted across time. We used indicator species analyses (indicspecies package; De Cáceres and Legendre, 2009) to determine ASVs associated with treatment groups at each time point. The indicator species analyses controls for type I error (false positives) by retesting the statistical significance of associations of significant ASVs (De Cáceres et al., 2010).

Since the abundance of bacteria within the Roseobacter group was of interest but they are not defined in the Silva taxonomic classifier, bacteria belonging to the Roseobacter clade were defined as having > 89% 16S rRNA sequence similarity (Buchan et al., 2005) to the only ASV listed as the genus Roseobacter in the dataset (ASV558). Lastly, a one-way ANOVA followed by Tukey’s post hoc test (stats R package, R Core Team, 2025) was used to investigate whether the total relative abundance of Roseobacter bacteria differed between treatments (total Roseobacter relative abundance ∼ treatment). Summarized Roseobacter data were confirmed to meet parametric ANOVA assumptions using a Q-Q plot to visualize heterogeneity of variance and the Shapiro-Wilks test for normality (*p* > 0.05).

## 4. Results

### 4.1 Bacterial Whole Genome Sequences – Roseobacter bacterium identified as *Celeribacter baekdonensis*

Shotgun sequencing generated 68 contigs (>100 bp), indicating an incomplete genome assembly (Supplementary Figure 2a); however, a complete genome was not required for our study. The bacterial isolate matched to *Celeribacter baekdonensis* (accession #: CP028475) with an ANI (average nucleotide identity) of 98.2% (genome alignment shown in Supplementary Figure 2b). The second-highest match between the probiotic isolate and all other reference genomes was to *Celeribacter neptunius* (79.3% ANI). The genome is relatively large for a bacterium (∼four million bps) and has high G/C nucleotide content (57.5%), which is characteristic of particle-associated Roseobacter (Luo et al., 2014). Phylogenetic relatedness between the *C. baekdonensis* probiotic 16S rRNA sequence and other Roseobacter genera found in the 16S rRNA dataset (described in section 4.7) is shown in Supplementary Figure 2c.

### 4.2 Larval survival – Roseobacter enrichment did not negatively impact larval development and improved larval survival amid high temperature exposure

Of the total 16 experimental tanks (four treatments replicated across four families), die-offs occurred in four tanks during larval rearing: three from the high temperature (HT) treatment and one from the control. Some variations in larval counts on 1 dpf are present (e.g., family 1, control), which may reflect variations in egg stocking densities rather than treatment effects (Figure 2a). During larval rearing, there were significant variations in survival across families and treatments. When oyster family was treated as a random factor, the HT treatment had significantly reduced larval survival (t = -2.95, *p* < 0.01), whereas the Roseobacter-enriched + HT treatment had no difference in survival (t = -0.28, *p* = 0.78) compared to the control (Figure 2b).

**Figure 2.**
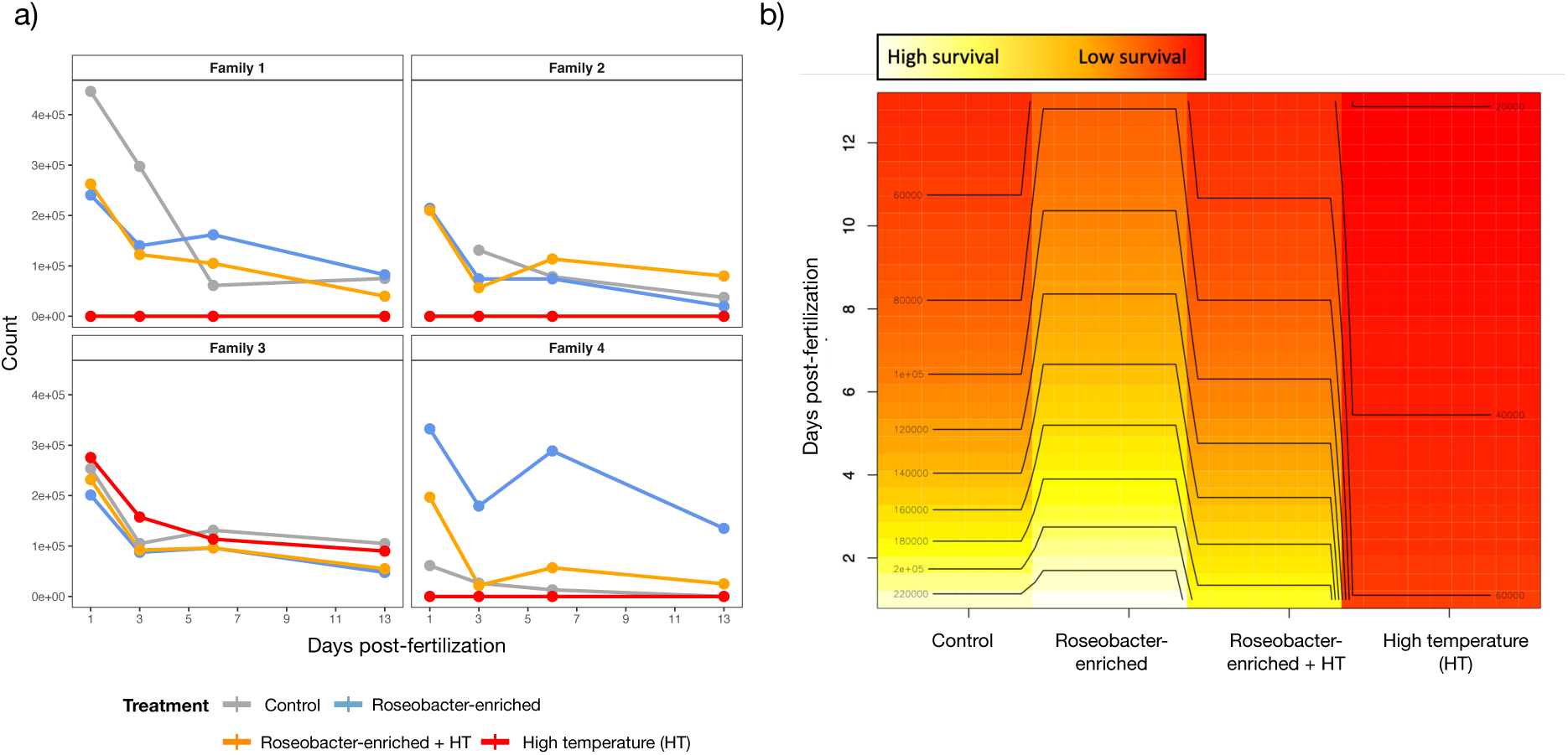
a) Averaged larval counts at each sampled day post-fertilization. b) Two-dimensional plot estimating larval survival (indicated by colour, high survival = white-yellow, low survival = red) across time (days post-fertilization, y-axis) between early life treatments, using a generalized additive mixed model with family as a random factor.

### 4.3 Larval (13 dpf) Vibrio aestuarianus + high temperature challenge – Survival varied between oyster families, but when family is statistically accounted for, larvae from the Roseobacter-enriched treatment have a 28% higher survival

Larval survival against *V. aestuarianus* differed between early-life treatments (chi-squared = 11.43(2), *p* < 0.01), family (chi-squared = 22.58(2), *p* < 0.01), and their interaction (chi-squared = 45.3(4), *p* < 0.01). Thereafter, treatment differences were examined within families, where the Roseobacter-enriched treatment had higher survival than the Roseobacter-enriched + HT treatment (z = -4.56, *p-adj* < 0.01) in family 2, and the Roseobacter-enriched + HT treatment had higher survival than the control in family 3 (z = 2.38, *p-adj* < 0.01) (Figure 3a). With family modelled as a random effect, larval survival against *V. aestuarianus* differed between early-life treatments (chi-squared = 8.95(2), *p* = 0.01), with the Roseobacter-enriched treatment having a mean 28% higher relative survival compared to the control (64% survival relative to 50% survival, z = 2.97, *p-adj* < 0.01). However, the Roseobacter-enriched + HT-control and Roseobacter-enriched-Roseobacter-enriched + HT were not significantly different (z = 1.33, *p-adj* = 0.548 and z = -1.79, *p-adj* = 0.220, respectively) (Figure 3b).

**Figure 3.**
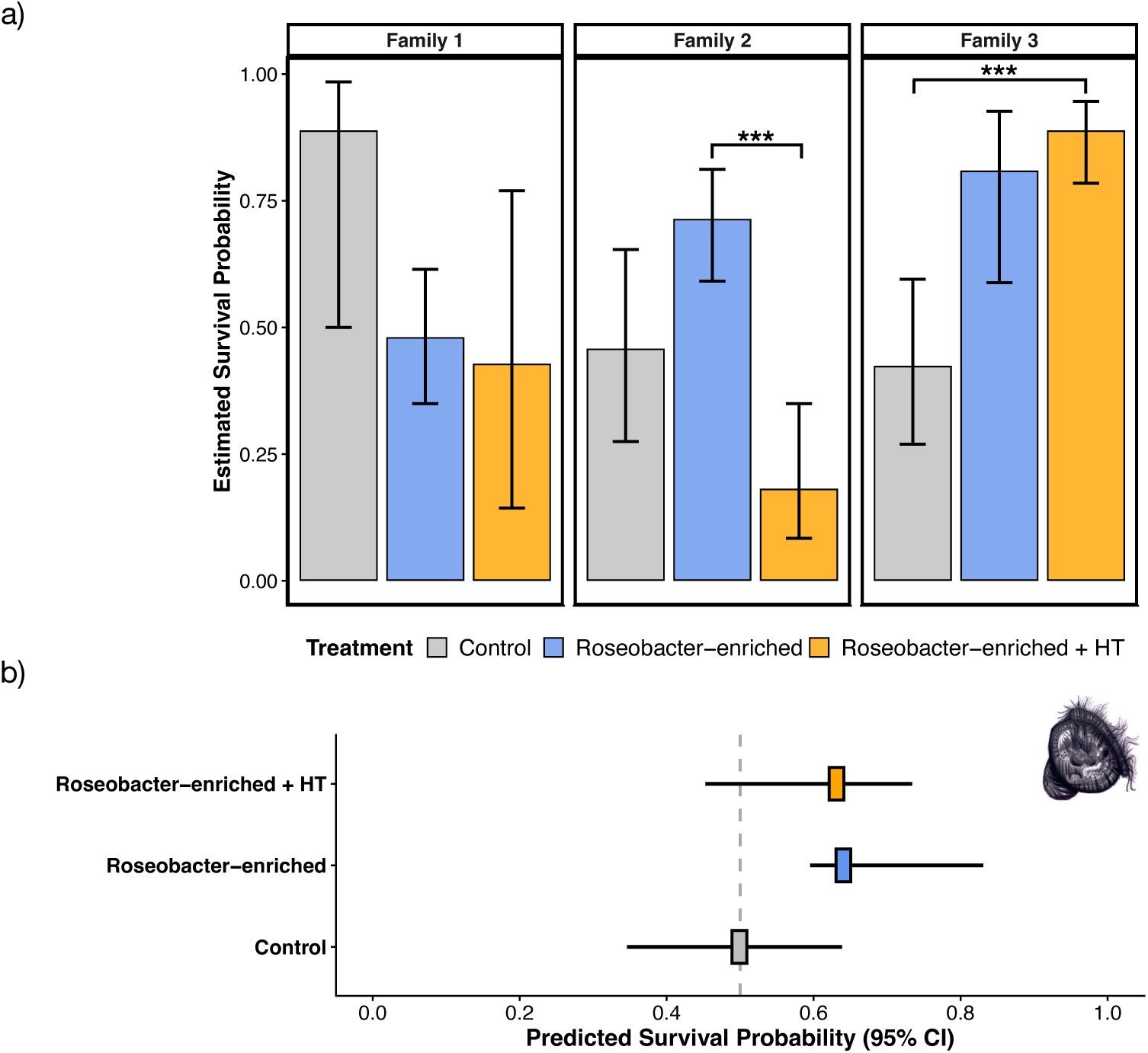
a) Estimated marginal mean survival probability following a *Vibrio aestuarianus* subsp*. francensis* and high temperature (24 °C) challenge with oyster larvae (13 days post-fertilization) originating from different early-life treatments [control (grey), Roseobacter-enriched (blue), Roseobacter-enriched + high temperature (HT) (orange)] and faceted by oyster family identity (Family 1, 2, and 3) (fixed effects model with the factors of treatment and family, AIC = 355). b) Predicted survival probability following a *Vibrio aestuarianus* subsp*. francensis* and high temperature (24 °C) challenge of larvae originating from different early-life treatments using a binomial generalized mixed model with oyster family as a random effect (AIC = 399.9).

### 4.4 Juvenile (90 dpf) Vibrio aestuarianus + high temperature challenge – Early Roseobacter enrichment improves oyster survival by ∼ 31%

Survival probability curves (fixed-effects model) indicated significant variation between treatments (*p* = 0.0065) (Figure 4a). When including family as a random effect (mixed effects model), Roseobacter-enriched and Roseobacter-enriched + HT treatments significantly reduced the hazard ratio (risk of death) by a mean of 30% (HR = 0.70, 95% CI = 0.53–0.93, *p* = 0.015) and 29% (HR = 0.71, 95% CI = 0.53–0.94, *p* = 0.017), respectively (*p* < 0.01) relative to the control (Figure 4b). When survival was compared within families, family 3 showed no significant treatment-related differences in survival when analyzed on its own (p = 0.29).

**Figure 4.**
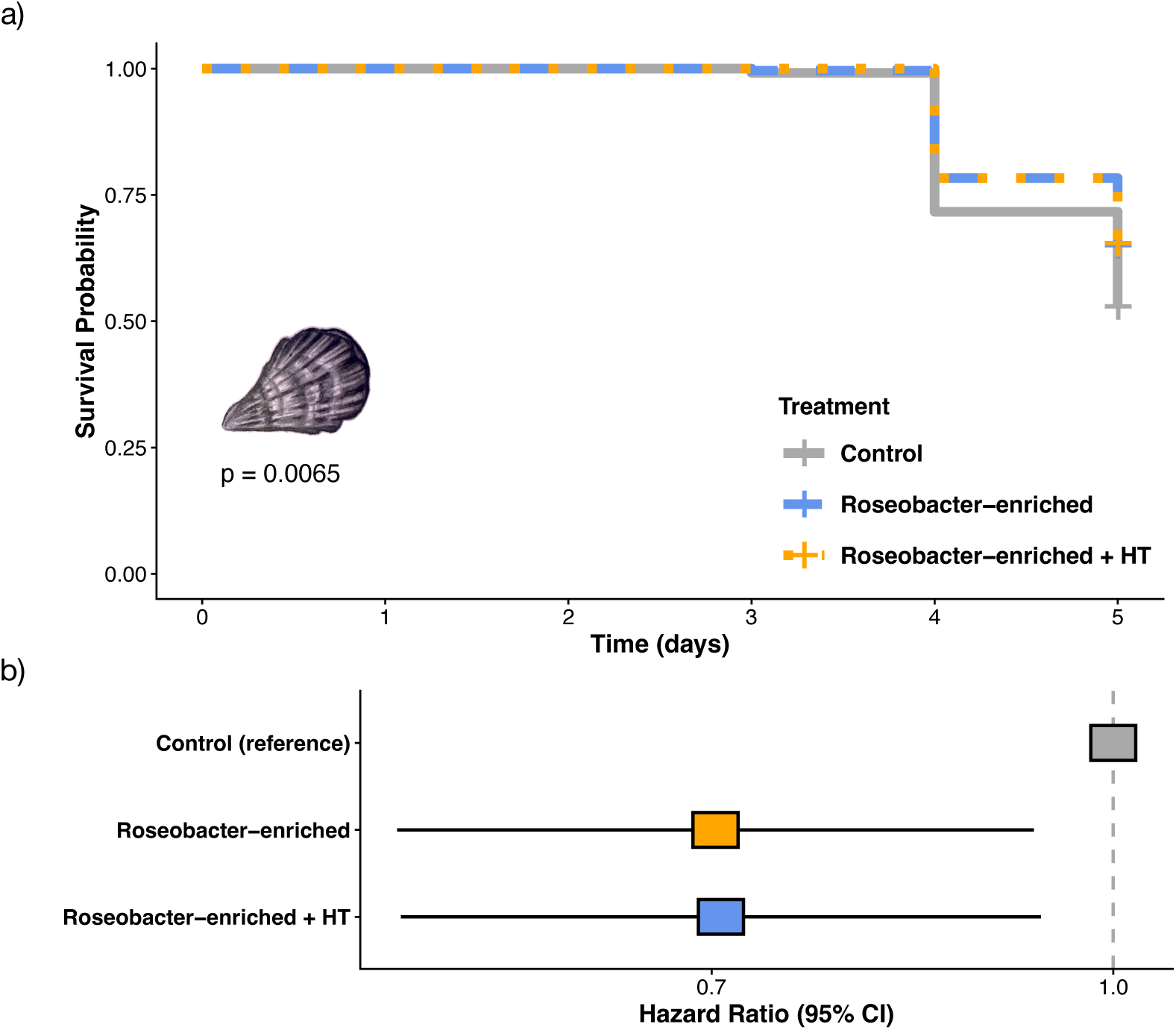
a) Survival probability over time (days following exposure to *Vibrio aestuarianus* pathogen) of Pacific oysters (90 days post-fertilization) originating from control (grey), Roseobacter-enriched (blue), or Roseobacter-enriched + HT (orange) treatments during the first 24 hours of life. b) Hazard ratio plot of a mixed-effects Cox model (family as a random effect) using survival data from the *Vibrio aestuarianus* + high temperature challenge (Integrated loglik AIC = 26.5). Oysters originating from the control treatment are set as the reference (hazard ratio = 1). ** *p <* 0.01.

### 4.5 Microbiome composition across time - Overall shifts in microbiome composition are driven by time and family, but not by early-life treatment

There were no differences in alpha diversity (diversity of ASVs) between treatments. When comparing alpha diversity across all time points, time had a significant effect on alpha diversity (Shannon, F=20.8(4), *p* < 0.01; inverse Simpson, X^2^=28.3(4), *p* < 0.01; Chao1, X^2^=25.3(4), *p* < 0.01). Post-hoc analyses indicated that the diversity of juvenile samples (90 dpf) was significantly higher than at all other time points (*p-adj* < 0.05 for all tests).

Changes in beta diversity (abundance of ASVs) are visualized as a PCoA plot, where overall differences between treatment groups diminish over time, and are not apparent by the juvenile stage (Figure 5). We used a PERMANOVA to test for overall differences in beta diversity between the factors of treatment, family, and their interaction at each time point (Supplementary Table 1). On 1 dpf, beta diversity was significantly different between treatments (F = 3.90(2), *p* < 0.01); however, there were no significant differences at any other time point. PERMANOVA indicated differences in beta diversity between families on 1, 3, 15, and 90 dpf (F = 2.63(2), *p* < 0.01; F = 4.16(2), *p* < 0.01; F = 0.382(2), *p* < 0.01, and F = 1.58(2), *p* = 0.01, respectively). Finally, at 90 dpf, there was a significant interaction effect between early life treatment and family (F = 1.98(4), *p <* 0.01).

**Figure 5.**
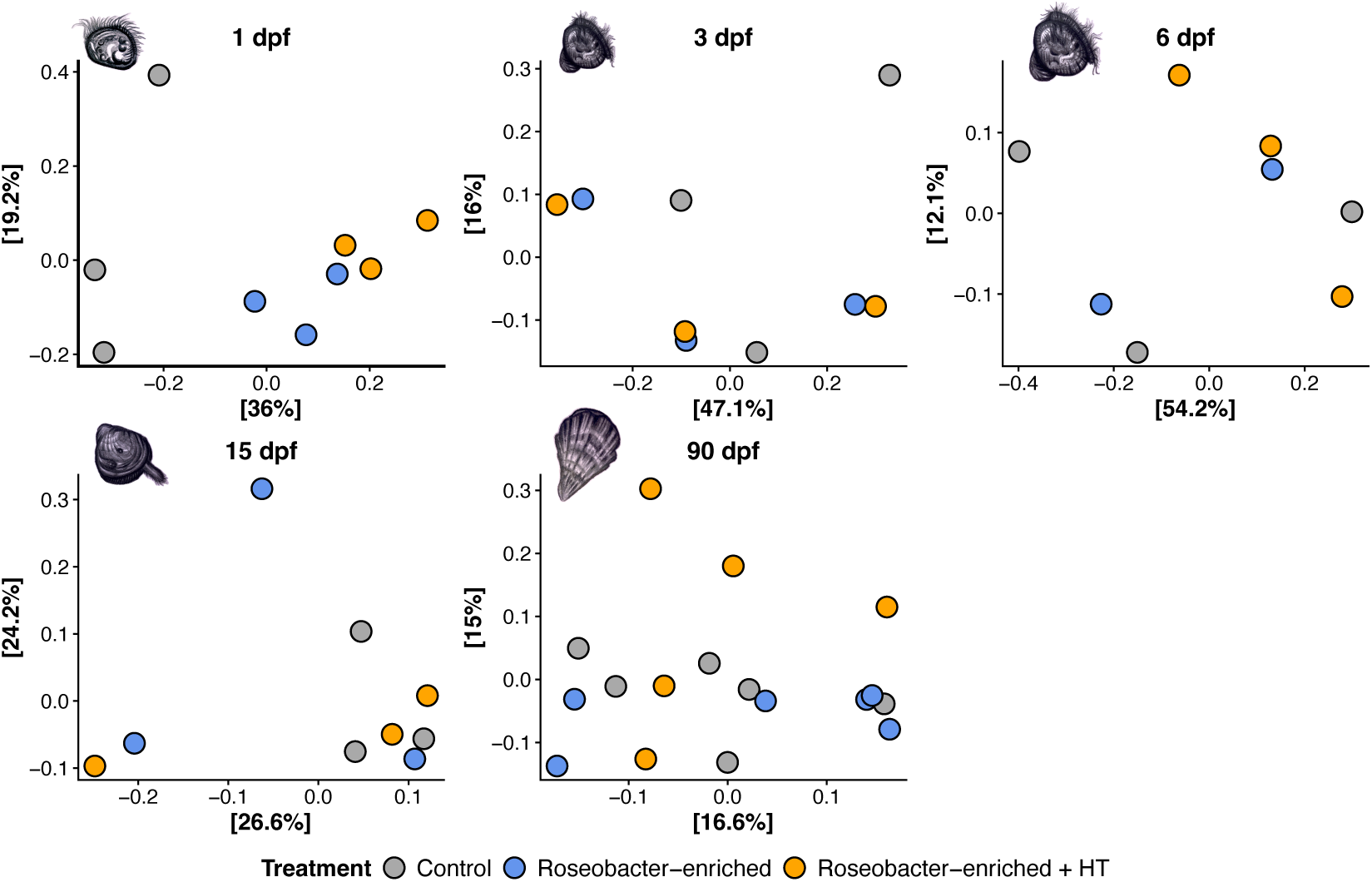
Principal component analysis (PCOa) plots of microbial diversity at each time-point [1 dpf (days post-fertilization), 3 dpf, 6 dpf, 15 dpf, and 90 dpf] using Brays-Curtis similarity matrices (999 permutations). Early life treatments are indicated by point colour where grey = control, blue = Roseobacter-enriched, and orange = Roseobacter-enriched + high temperature (HT). *p*-values indicate significant effects of treatment on beta-diversity at each time point, where ns = non-significant.

### 4.6 The core microbiome – Few core microbes are present across all time points, but the majority are Roseobacter bacteria

Analysis of each treatment’s core microbiome (90% prevalence threshold) revealed that only five core ASVs were present across all time points. The majority of core ASVs are Roseobacter bacteria (4/5 core ASVs), with most belonging to the *Phaeobacter* genus (3/5 core ASVs). Most core ASVs were shared between treatments except ASV8 (*Alteromonas*) in the control, and ASV11 (unknown Roseobacter) in the Roseobacter-enriched treatment. Additionally, the control lacked a core ASV shared between the Roseobacter-enriched and Roseobacter-enriched + HT treatments (ASV3; *Phaeobacter*) (Figure 6, Venn diagram). The abundance of all core ASVs through time is shown in Figure 6.

**Figure 6.**
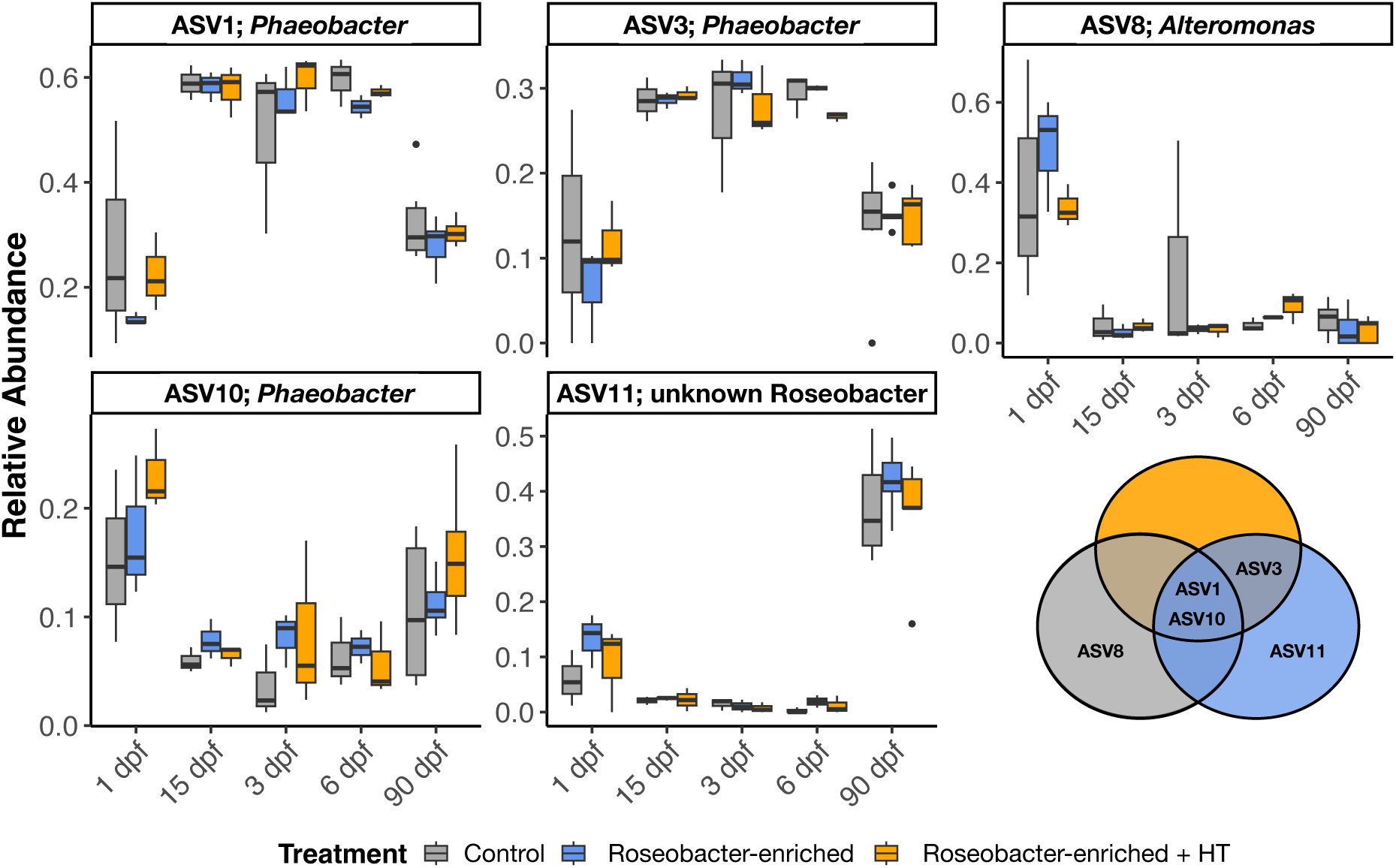
Relative abundance of core bacterial members (Amplicon Sequence Variants, ASVs) shared between treatments (shown in Venn diagram) across all time points (dpf = days post-fertilization). Grey = control, blue = Roseobacter-enriched, and orange = Roseobacter-enriched + high temperature (HT).

### 4.7 Roseobacter abundance – Although the abundance of C. baekdonensis diminishes over time, Roseobacter enrichment results in an increased abundance of specific and total Roseobacter bacteria at 90 dpf

The *C. baekdonensis* bacterium did not remain associated with *C. gigas* over time. Following Roseobacter enrichment at 1 dpf, the abundance of *C. baekdonensis* declined over time, reaching zero by the juvenile stage (Figure 7a). At the juvenile stage, several Roseobacter ASVs were associated with the Roseobacter-enriched and/or Roseobacter-enriched + HT treatment groups (full results in Supplementary Table 2). The ASVs 88 and 178 were significantly associated with the Roseobacter-enriched and Roseobacter-enriched + HT treatments (Figure 7c). The ASVs 88 and 178 have a 97% sequence similarity, and the top NCBI nBLAST matches were to sequences of the *Loktanella* genus. Additionally, several Roseobacter ASVs (*Sulfitobacter* genus and unknown Roseobacter) were significantly associated with only the Roseobacter-enriched treatment (Figure 7d). When comparing the total relative abundance of Roseobacter bacteria across treatment, family, or their interaction, only treatment had a significant effect (F = 6.13(2), *p* = 0.024). Tukey’s post-hoc analysis indicated that the Roseobacter-enriched treatment had a significantly higher abundance of Roseobacter bacteria compared to the control (*p-adj* = 0.033) but not the Roseobacter-enriched + HT treatment (*p-adj* = 0.096); however, the Roseobacter-enriched and Roseobacter-enriched + HT treatments were not different (*p-adj* = 0.897).

**Figure 7.**
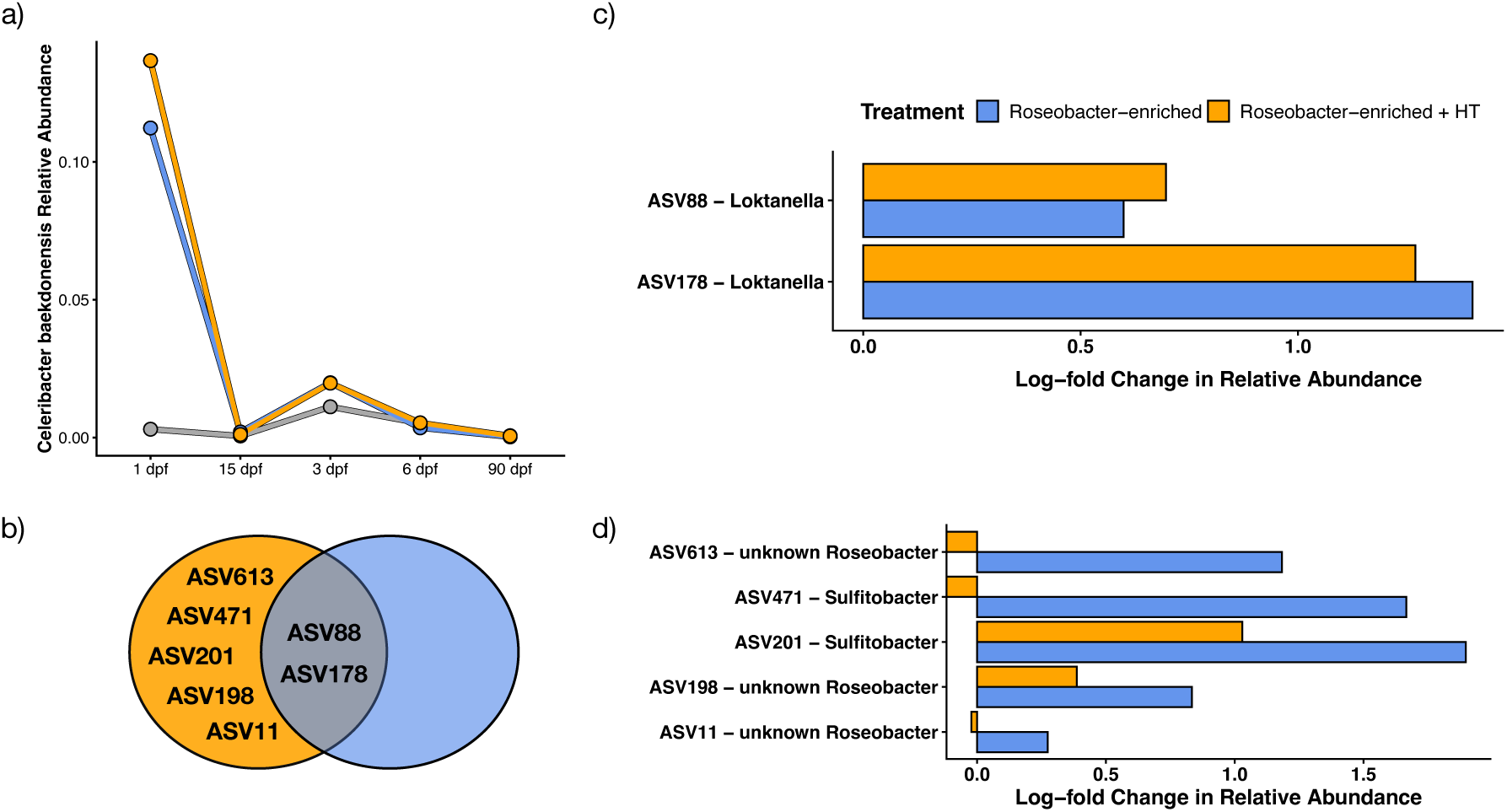
a) Relative abundance of the probiotic bacteria over time. Grey = control, blue = Roseobacter-enriched, orange = Roseobacter-enriched + HT. b) Venn diagram of Roseobacter amplicon sequence variants (ASV) significantly associated with the Roseobacter-enriched and/or Roseobacter-enriched + high temperature (HT) treatment groups. Log-fold change in the relative abundance of Roseobacter ASVs significantly associated with c) both the Roseobacter-enriched and Roseobacter-enriched + HT treatments or d) only the Roseobacter-enriched treatment, relative to the control. Asterix (*) indicates *p* < 0.05 and double Asterix (**) indicates *p* < 0.01. <unknown Roseobacter= indicates ASVs with an unknown genus that have a >89% match to a Roseobacter in the dataset.

## 5. Discussion

Microbial exposure during early development is increasingly recognized as a critical window in which environmental conditions can shape long-term host–microbe relationships and, ultimately, host health. Here, we tested this “microbial education” framework in *C. gigas* by manipulating the microbial and temperature environment during the first day of life and then tracking consequences for development, microbiome composition, and survival against disease across later life stages. Our findings indicate that brief but early enrichment with a Roseobacter bacteria (*C. baekdonensis*) can have enduring effects on host outcomes, including larval survival, altered microbiome assembly, and improved robustness to pathogen- and temperature-associated stress across multiple life stages (larval and juvenile). Notably, these benefits emerged even though the supplemented bacterium itself was not maintained within the host’s microbiome, suggesting that early exposure may act by steering subsequent microbial community trajectories and/or host immune function rather than by simple persistence of the supplemented bacteria. At the same time, large family-level differences underscore that host genetic background can modulate the impacts of early microbial exposure. Although speculative, the high abundance of Roseobacter bacteria later in life could be mediated by alterations in the host immune system, such as the production of antimicrobial peptides (AMPs) (Fallet et al., 2022), and these changes could be imprinted into innate immune memory by unknown cellular mechanisms. Together, these results support microbial education hypotheses, where an organism’s first microbial exposure shapes long-term host-microbe interactions with both pathogens and potentially protective bacteria.

### 5.1 Exposure to high temperature reduced larval survival, but the addition of Roseobacter bacteria resulted in a “microbial rescue effect.”

Early-life Roseobacter enrichment over the first 24 hours showed no adverse effects on larval growth or survival, supporting its potential as a microbial education strategy in *C. gigas*. Moreover, Roseobacter enrichment improved larval survival against the negative effects of high temperature exposure, demonstrating a “microbial rescue effect” from Roseobacter enrichment (Lehtinen, 2023). The observed positive-to-neutral effects of Roseobacter enrichment on larval survival contrast with other studies reporting moderate-to-severe adverse effects on larval development (Brown, 1973; Douillet and Langdon, 1993; Abasolo-Pacheco et al., 2017; Datan et al., 2024). Heat stress can reduce microbial diversity and stability in bivalves (Lokmer et al., 2015; Li et al., 2019) and the immune activity of *C. gigas* (e.g., Malham et al., 2009). Exposure to *C. baekdonensis* may have bolstered an immune response that would otherwise be suppressed during high temperature exposure, improving survival during larval development. Although research on this topic in oysters is scant, previous studies on Kurama shrimp (*Marsupenaeus japonicus*) have shown that adding *Clostridium butyricum* probiotics increases intestinal antioxidant capacity, thereby improving heat tolerance (Duan et al., 2017). This raises the question of whether microbial supplementation could moderate temperature stress at later life stages. Roseobacter supplementation has been used to mitigate coral mortality (Li et al., 2023), coral bleaching (Rosado et al., 2019), and to improve coral skeleton structure during heat stress (30 - 31 °C) and/or *Vibrio coralliilyticus* exposure (Moradi et al., 2023). However, the use of microbial supplementation to mitigate temperature stress in other sessile marine organisms remains less researched. Here, we present *C. baekdonensis* as a candidate bacterium to explore these concepts in *C. gigas*.

### 5.2 Application of Roseobacter enrichment to improve survival against Vibrio aestuarianus

Roseobacter enrichment during the first 24 hours improved survival against *V. aestuarianus* infection and high temperatures at 13 dpf by 28% (Roseobacter-enriched treatment only) and 90 dpf by ∼ 30% (Roseobacter-enriched and Roseobacter-enriched + HT treatment). Similarly, Datan et al., found that microbial enrichment of seawater from healthy donor oysters from 3 hours post-fertilization to 14 dpf improved oyster survival against *V. aestuarianus* one year later by ∼ 28% (2024). This suggests that prolonged microbial enrichment may not improve long-term benefits, and that early exposure (i.e., the first 24 hours) is key. The absence of compounding beneficial effects from continuous microbial exposure emphasizes the temporal limitations of microbial education processes. Datan et al., also found that several microbial mixtures (10^4^ cells·mL^-1^·day^-1^) from 3 hours post-fertilization – 14 dpf or 7 – 14 dpf did not improve survival against *V. aestuarianus* one year later, suggesting that the protective effects of microbial education may be taxon dependent (Datan et al., 2024). Finding bacterial taxa that consistently increase long-term disease resilience, or, in other words, the best “teachers” for microbial education processes, will be an important next step in this research.

The beneficial effects of Roseobacter-enrichment across life stages were not found in all families. At 13 and 90 dpf, families 1 and 3, respectively, showed no differences in survival during *V. aestuarianus* and high temperature challenges between early-life treatments when examined independently. Family-level variation in survival underscores the need to replicate early-life treatments across different genetic backgrounds when assessing the long-term effects of microbial education. These results highlight that changes in host processes following Roseobacter enrichment may be complex and interact with unknown genetic factors.

### 5.3 Increased survival against V. aestuarianus and long-term microbiome shifts in the Roseobacter community following Roseobacter enrichment

Our results suggest that microbial education processes may be involved in a host’s long-term defence against pathogens and in the selection of its microbiota. There are several cellular mechanisms which may underlie these processes. One likely explanation is that the beneficial effects of Roseobacter enrichment on survival observed in this study may be mediated by shaping the innate immune system to mount more effective defences against pathogens and to select for Roseobacter that may protect against pathogens. Early exposure to microbial-enriched seawater in the first 24 hours of life can alter *C. gigas* gene expression, including the upregulation of host-derived AMPs (Fallet et al., 2018). A possible mechanism for the increased abundance of Roseobacter bacteria following Roseobacter enrichment is that early microbial exposure induces immune changes in the host, such as increased AMP expression. Increased production of host-derived AMPs may exert selective pressure for AMP-tolerant bacteria in the environment (Destoumieux-Garzón et al., 2024). Such AMP-resistant bacteria may produce their own AMPs, bolstering the host’s disease robustness (Desriac et al., 2014). Bacterial resistance to many host-derived AMPs requires the ability to remove protons (proton efflux), which is the same cellular strategy Roseobacter use to tolerate TDA (Joo et al., 2016; Wilson et al., 2016). The chemical selection of microbiota through host-derived AMPs is widespread across animals, including marine invertebrates, and plays a role in defence against pathogens and in the selection of symbionts (Destoumieux-Garzón et al., 2016).

Changes in survival against *V. aestuarianus* occurred with key shifts in the Roseobacter community. The total relative abundance of all Roseobacter bacteria was higher in juveniles from the Roseobacter-enriched treatment compared to the control. Juveniles from the Roseobacter-enriched + HT treatment had intermediate Roseobacter abundance. Although there were no differences in Roseobacter abundance at the larval stages (3 - 15 dpf), we did find that certain Roseobacter (*Phaeobacter* and unknown Roseobacter) were persistent core microbes across developmental stages, with an additional Roseobacter belonging to the *Phaeobacter* genus being a core member of the Roseobacter-enriched treatments. Members of the *Phaeobacter* genus are known to produce TDA (Bruhn et al., 2007; Geng and Belas, 2008) and inhibit Vibrio pathogens (Ruiz-Ponte et al., 1999). Other Roseobacter genera associated with the Roseobacter-enriched treatments at 90 dpf include *Sulfitobacter*, *Loktanella*, and three Roseobacter ASVs not resolved to the genus level (Figure 7). These Roseobacter bacteria could represent important microbes for *C. gigas* health and protection against *Vibrio aestuarianus* pathogens.

Overall, research on the beneficial effects of *Sulfitobacter* and *Loktanella* on bivalve survival against *Vibrio* pathogens, including the production of TDA or other antimicrobial compounds, is limited. *Sulfitobacter* has been shown to inhibit *Vibrio anguillarum* growth by producing unknown antibacterial compounds and competing for phytoplankton-derived nutrients (Sharifah and Eguchi, 2011). However, a study on two *Sulfitobacter* species tested (1921; TM1040 and EE36) found that neither produces TDA in response to TDA-chemical signalling (autoinduction pathway) based on reporter gene assays (Geng and Belas, 2010), suggesting *Sulfitobacter* may produce other, unidentified antimicrobial compounds. Although few studies have investigated the antibacterial activity of the genus *Loktanella* (also classified in the *Yoonia* genus; Wirth et al., 2018), *Loktanella maritima* crude extracts have been shown to inhibit *V. anguillarum*, an American lobster (*Homarus americanus*) pathogen, when cocultured with other bacteria (Ranson, 2020). Genotypic analysis of *L. maritima* revealed the presence of gene clusters involved in the production of antibacterial compounds (Ranson et al., 2018). Notably, the abundance of *Sulfitobacter* and *Loktanella* is enriched in DMSP-concentrated environments (Mou et al., 2005), a sulphur compound that can act as a signalling molecule and a sulphur source for TDA production (Geng and Belas, 2010), suggesting that further research should be conducted to confirm antimicrobial activities of *Sulfitobacter* and *Loktanella*.

## 6. Conclusion

Microbial exposure during early development appears to be a meaningful lever that shapes host–microbe relationships and later disease outcomes. In this study, early-life enrichment with Roseobacter bacteria supports the broader concept of microbial education in marine invertebrates, suggesting that brief encounters with beneficial taxa early in life can influence subsequent microbiome assembly and host survival against pathogen and warming-associated stress. We propose that enhanced survival during disease challenges, along with long-term microbiome changes, may be mediated by alterations in the host immune response and Roseobacter abundance. Family-level variations in survival indicate that host genetic background may moderate these effects, emphasizing that microbial interventions will likely interact with family-level differences in immune and microbiome trajectories. While our experimental design necessarily reflects the logistical constraints of hatchery-scale experiments, our findings highlight both the promise and the complexity of using targeted microbes to steer long-term health outcomes. Overall, integrating microbial education into bivalve rearing offers a practical, ecologically grounded approach for mitigating climate-amplified disease risk in aquaculture, conservation, and restoration. Future studies should focus on identifying the most effective microbial “teaching” consortia, testing broader genetic and environmental contexts, and uncovering the immune and molecular mechanisms that underpin persistent protective effects.

## 7. Supplementary Materials

**Supplementary Figure 1.**
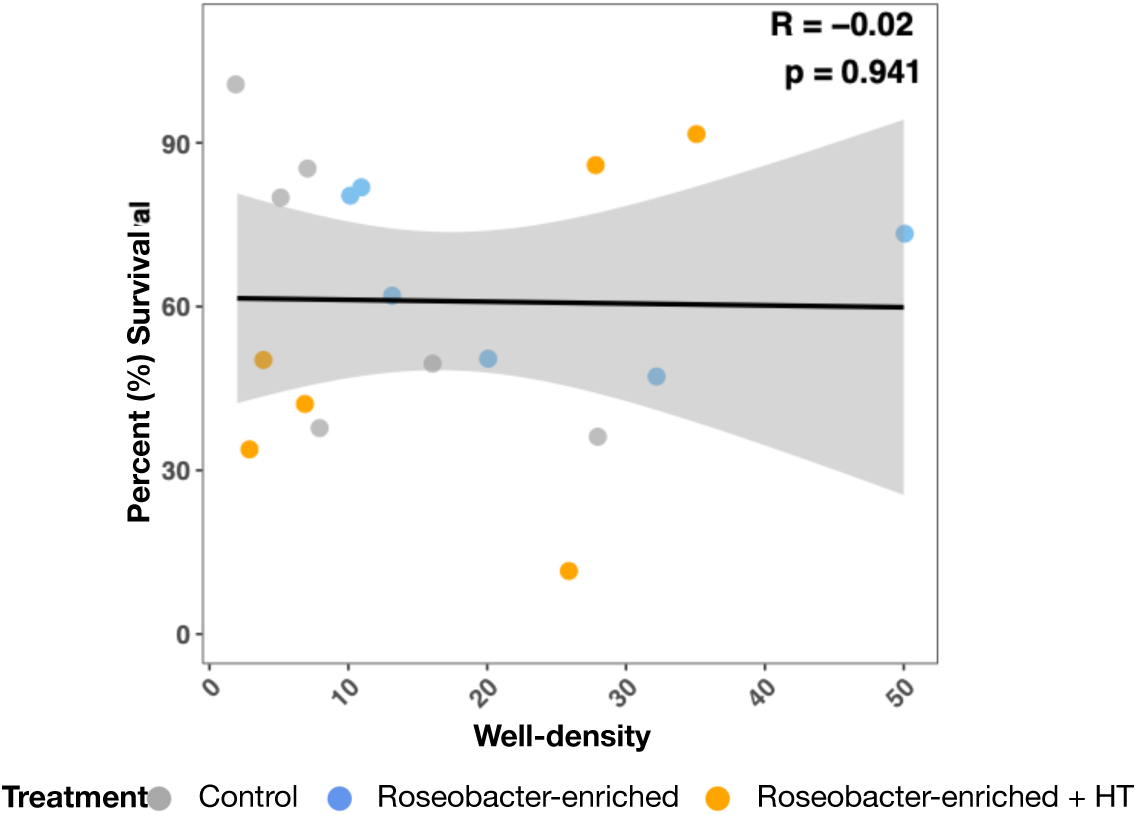
Correlation plot and Pearson correlation test between larval (13 days post fertilization) percent (%) survival against *Vibrio aestuarianus* subsp. *francensis* + high temperature (24 °C) and well density (number of larvae per well) during the disease challenge.

**Supplementary Figure 2.**
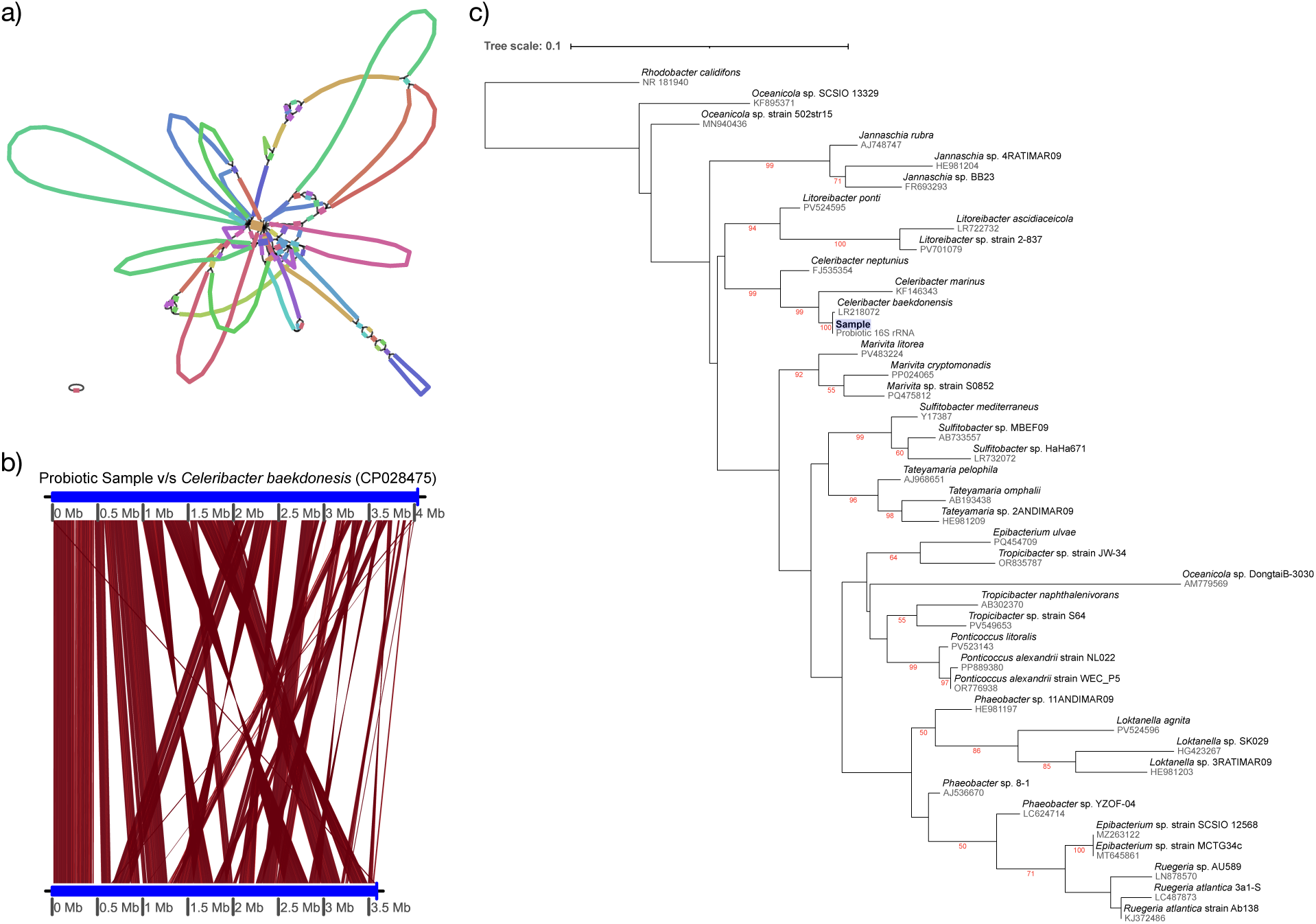
a) Visualization of contigs representing final genome assembly. b) Average nucleotide identification compared between *Celeribacter baekdonensis* (accession: #CP028475) and assembly. c) Maximum likelihood phylogenetic reconstruction of 16S rRNA genes within the Roseobacter Group.

**Supplementary Table 1.**
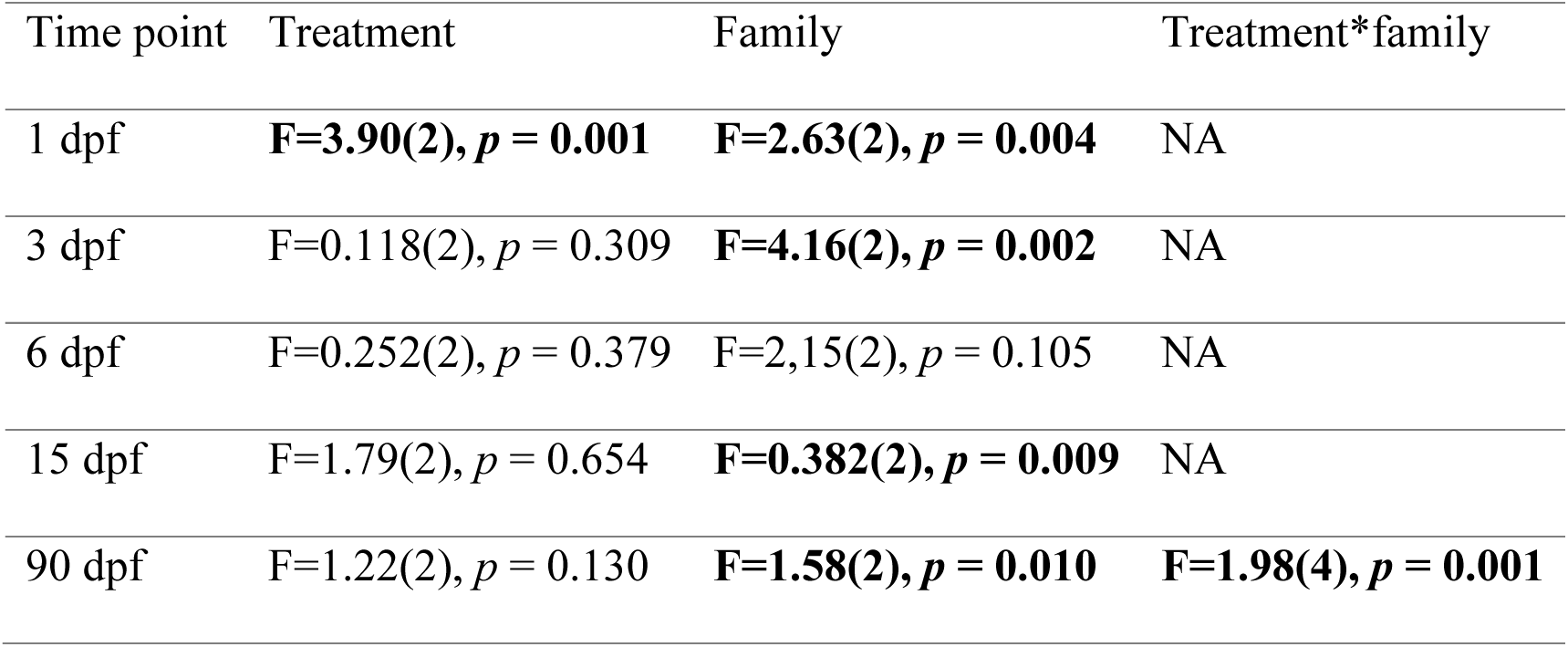
Results of Permutational Multivariate Analysis of Variance (PERMANOVA) tests with the factors treatment, family, and their interaction (Treatment*Family) at each time-point to detect differences in microbiome composition (beta-diversity). At some time points it was not possible to test for interactive effects of treatment and family due to low sample size (NA = not available). Significant results (*p* < 0.05) are in bold. Dpf = days post-fertilization.

**Supplementary Table 2.**
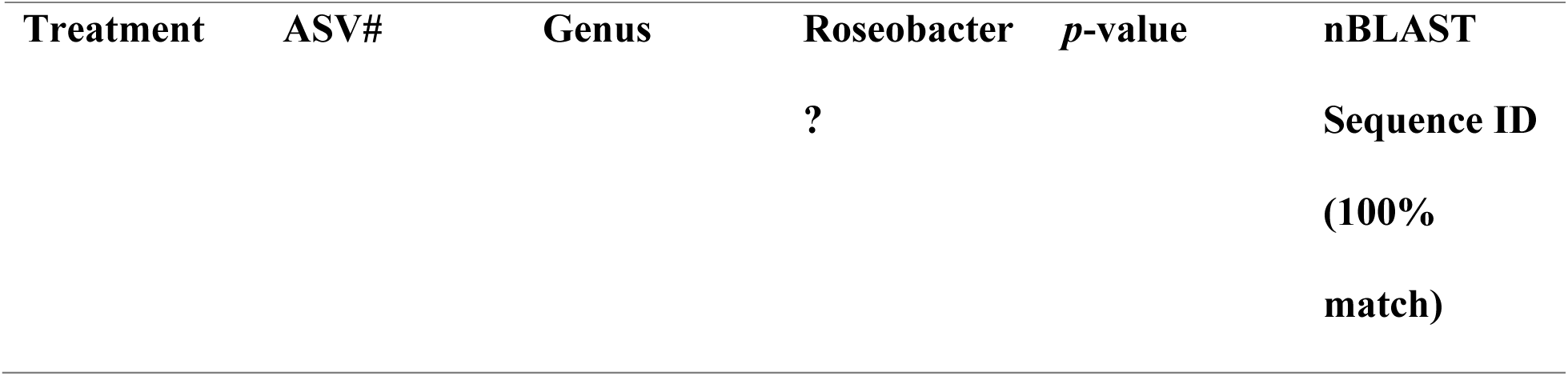
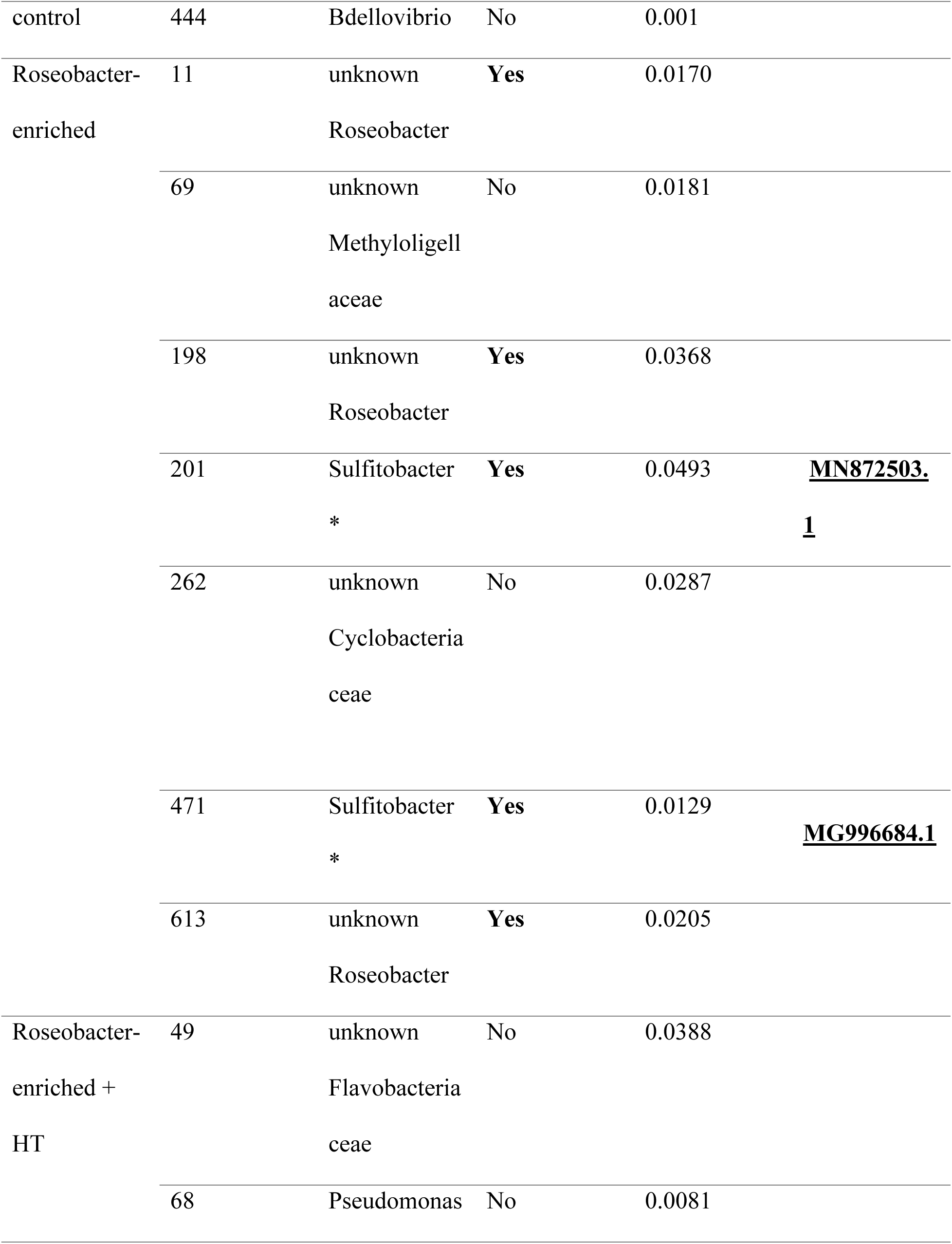
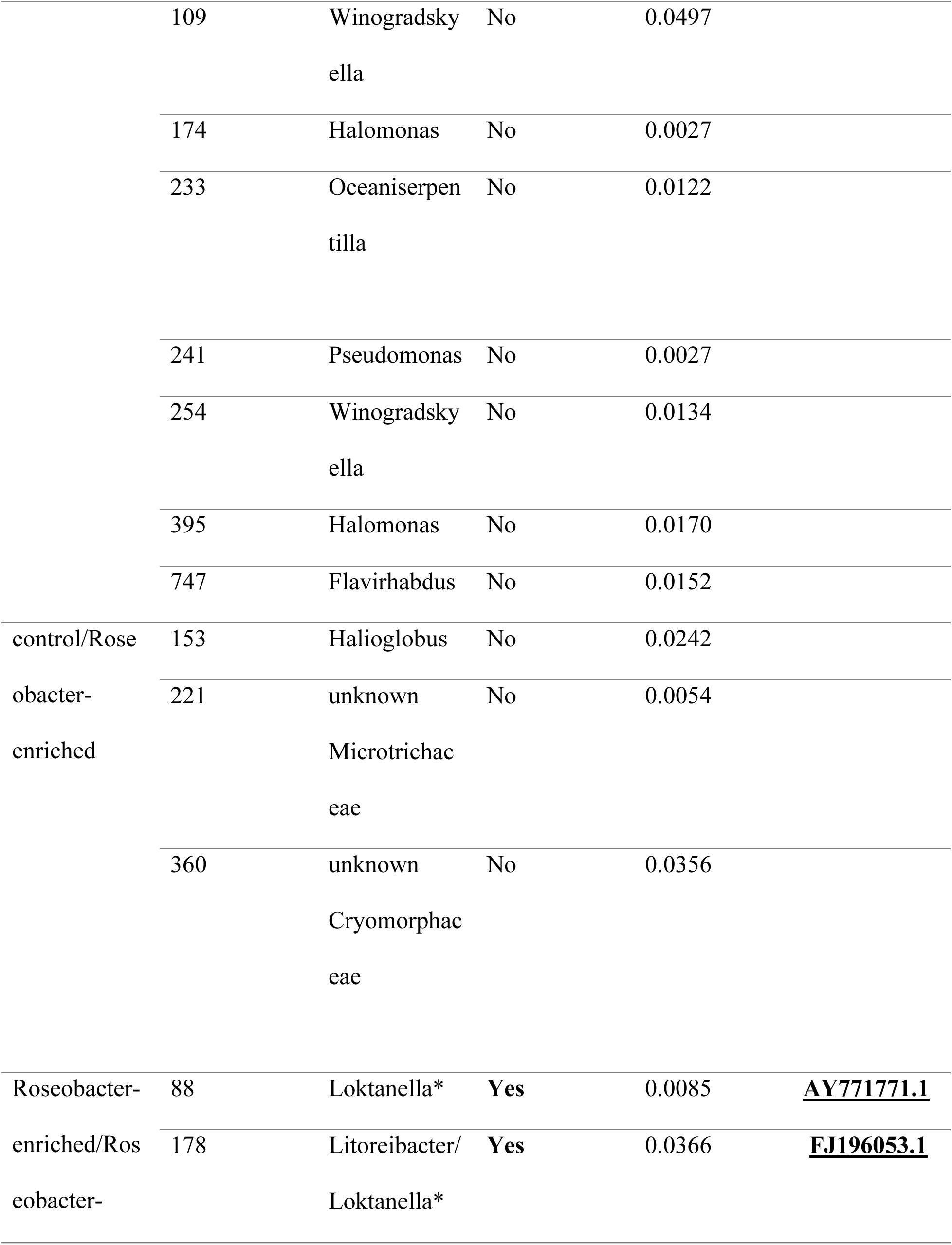
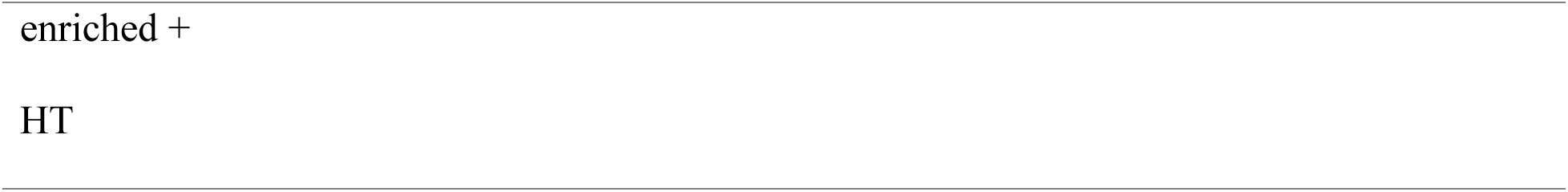
Indicator species analysis results. Significant Amplicon Sequence Variants (ASVs) (*p* < 0.05) indicate the association between the ASV abundance with one or more treatment groups. ASVs are defined as a Roseobacter if they are one of the described genera within the Roseobacter group and/or if the 16S rRNA sequence matches > 89% with the Roseobacter ASV within the dataset. HT = High temperature. Asterix indicates that the NCBI nBLAST database was used to search for sequence ID matches (100%).

## 8. Data availability

Raw 16S rRNA data is available at Harvard Dataverse: https://doi.org/10.7910/DVN/LRN7DL

All code is provided at the GitHub repository: https://github.com/MarissaLag/MU42022_Code

## 9. Acknowledgements

The authors acknowledge funding from the Canada Research Chair Program (CRC, #950–231856) and the Natural Sciences and Engineering Research Council (NSERC, #2018–06761) to T.J.G, and NSERC Canadian Graduate Scholarship –Doctoral program (NSERC, #600069-2025) to M.W.L. The authors thank the Deep Bay Marine Field Station staff, Carl Butterworth, Sarah Leduc, Daniel Roth, Chloe McLaughlin, and Megan Lebeuf, who made the hatchery experiment possible. We thank Bri Cooper for illustrating the larval images.

## References

Abasolo-Pacheco, F., Campa-Córdova, Á.I., Mazón-Suástegui, J.M., Tovar-Ramírez, D., Araya, R. and Saucedo, P.E., 2017. Enhancing growth and resistance to *Vibrio alginolyticus* disease in catarina scallop (*Argopecten ventricosus*) with Bacillus and Lactobacillus probiotic strains during early development. Aquaculture Research, 48(9), pp.4597–4607.

Andrews, S., 2010. FastQC: a quality control tool for high-throughput sequence data. Available online at: http://www.bioinformatics.babraham.ac.uk/projects/fastqc/ (v0.12.1).

Balcázar, J.L., Rojas-Luna, T. and Cunningham, D.P., 2007. Effect of the addition of four potential probiotic strains on the survival of pacific white shrimp (*Litopenaeus vannamei*) following immersion challenge with *Vibrio parahaemolyticus*. Journal of invertebrate pathology, 96(2), pp.147–150.

Bates D., Maechler M., Bolker B., and Walker S., 2015. Fitting Linear Mixed-Effects Models Using lme4. Journal of Statistical Software, 67(1), pp. 1–48. doi:10.18637/jss.v067.i01.

Bekkering, S., Domínguez-Andrés, J., Joosten, L.A., Riksen, N.P. and Netea, M.G., 2021. Trained immunity: reprogramming innate immunity in health and disease. Annual review of immunology, 39, pp. 667–693.

Bolger, A. M., Lohse, M., & Usadel, B., 2014. Trimmomatic: A flexible trimmer for Illumina sequence data. Bioinformatics, 30(15), pp.2114–2120. 10.1093/bioinformatics/btu170.

Brinkhoff, T., Bach, G., Heidorn, T., Liang, L., Schlingloff, A. and Simon, M., 2004. Antibiotic production by a Roseobacter clade-affiliated species from the German Wadden Sea and its antagonistic effects on indigenous isolates. Applied and environmental microbiology, 70(4), pp.2560–2565.

Brown, C., 1973. The effects of some selected bacteria on embryos and larvae of the American oyster, *Crassostrea virginica*. Journal of Invertebrate Pathology, 21(3), pp.215–223.

Bruhn, J.B., Nielsen, K.F., Hjelm, M., Hansen, M., Bresciani, J., Schulz, S. and Gram, L., 2005. Ecology, inhibitory activity, and morphogenesis of a marine antagonistic bacterium belonging to the Roseobacter clade. Applied and environmental microbiology, 71(11), pp.7263–7270.

Bruhn, J.B., Gram, L. and Belas, R., 2007. Production of antibacterial compounds and biofilm formation by *Roseobacter* species are influenced by culture conditions. Applied and environmental microbiology, 73(2), pp. 442–450.

Buchan, A., González, J.M. and Moran, M.A., 2005. Overview of the marine Roseobacter lineage. Applied and environmental microbiology, 71(10), pp.5665–5677.

Burge, C.A. and Hershberger, P.K., 2020. Climate change can drive marine diseases. Oxford: Oxford University Press, pp. 83–94.

Callahan, B.J., Sankaran, K., Fukuyama, J.A., McMurdie, P.J. and Holmes, S.P., 2016. Bioconductor workflow for microbiome data analysis: from raw reads to community analyses. F1000Research, 5, pp.1492.

Cousson, A., Cheutin, M.C., Rieuvilleneuve, F., Roques, C., Pouzadoux, J., Voron, F., Mas, S., Violette, H., Hatey, E., Got, P. and Gros, R., 2025. Synergistic pathogenicity between Herpesvirus and *Vibrio aestuarianus* in killing juvenile oysters during spring in natural conditions. Science of The Total Environment, 1003, pp.180675.

Cowan, M.W., Pearce, C.M., Finston, T., Meyer, G.R., Marshall, R., Evans, W., Sutherland, T.F. and De La Bastide, P.Y., 2023. Role of the *Vibrio* community, reproductive effort, and environmental parameters in intertidal Pacific oyster summer mortality in British Columbia, Canada. Aquaculture, 565, pp.739094.

Cowan, M.W., Pearce, C.M., Green, T.J., Finston, T., Meyer, G.R., McAmmond, B., Van Hamme, J.D., Bottos, E.M., Marshall, R., Evans, W. and Sutherland, T.F., 2024. Abundance of *Vibrio aestuarianus*, water temperature, and stocking density are associated with summer mortality of Pacific oysters in suspended culture. Aquaculture International, 32(4), pp. 5045–5066.

Coyle, N.M., O’Toole, C., Thomas, J.C., Ryder, D., Feil, E.J., Geary, M., Bean, T.P., Joseph, A.W., Waine, A., Cheslett, D. and Verner-Jeffreys, D.W., 2023. *Vibrio aestuarianus* clade A and clade B isolates are associated with Pacific oyster (*Magallana gigas*) disease outbreaks across Ireland. Microbial Genomics, 9(8).

Dantan, L., Toulza, E., Petton, B., Montagnani, C., Dégremont, L., Morga, B., Fleury, Y., Mitta, G., Gueguen, Y., Vidal-Dupiol, J. and Cosseau, C., 2024. Microbial education for marine invertebrate disease prevention in aquaculture. Reviews in Aquaculture, 16(3), pp. 1229–1243.

Dantan, L., Carcassonne, P., Degrémont, L., Morga, B., Travers, M.A., Petton, B., Mege, M., Maurouard, E., Allienne, J.F., Courtay, G. and Romatif, O., 2024. Microbial education plays a crucial role in harnessing the beneficial properties of microbiota for infectious disease protection in *Crassostrea gigas*. Scientific Reports, 14(1), p.26914.

Darriba, D., Posada, D., Kozlov, A. M., Stamatakis, A., Morel, B., & Flouri, T., 2020. ModelTest-NG: A New and Scalable Tool for the Selection of DNA and Protein Evolutionary Models. Molecular Biology and Evolution, 37(1), pp. 291–294. 10.1093/molbev/msz189

De Cáceres, M., Legendre, P. and Moretti, M., 2010. Improving indicator species analysis by combining groups of sites. Oikos, 119(10), pp.1674–1684.

de Lorgeril, J., Escoubas, J.M., Loubiere, V., Pernet, F., Le Gall, P., Vergnes, A., Aujoulat, F., Jeannot, J.L., Jumas-Bilak, E., Got, P. and Gueguen, Y., 2018. Inefficient immune response is associated with microbial permissiveness in juvenile oysters affected by mass mortalities on field. Fish & shellfish immunology, 77, pp. 156–163.

Desriac, F., Le Chevalier, P., Brillet, B., Leguerinel, I., Thuillier, B., Paillard, C. and Fleury, Y., 2014. Exploring the hologenome concept in marine bivalvia: haemolymph microbiota as a pertinent source of probiotics for aquaculture. FEMS microbiology letters, 350(1), pp. 107–116.

Destoumieux-Garzón, D., Rosa, R.D., Schmitt, P., Barreto, C., Vidal-Dupiol, J., Mitta, G., Gueguen, Y. and Bachere, E., 2016. Antimicrobial peptides in marine invertebrate health and disease. Philosophical Transactions of the Royal Society B: Biological Sciences, 371(1695).

Destoumieux-Garzón, D., Montagnani, C., Dantan, L., Nicolas, N.D.S., Travers, M.A., Duperret, L., Charrière, G.M., Toulza, E., Mitta, G., Cosseau, C. and Escoubas, J.M., 2024. Cross-talk and mutual shaping between the immune system and the microbiota during an oyster’s life. Philosophical Transactions of the Royal Society B, 379(1901).

Dinno A., 2024. dunn.test: Dunn’s Test of Multiple Comparisons Using Rank Sums. R package version 1.3.6, https://CRAN.R-project.org/package=dunn.test.

Douillet, P. and Langdon, C.J., 1993. Effects of marine bacteria on the culture of axenic oyster Crassostrea gigas (Thunberg) larvae. The Biological Bulletin, 184(1), pp. 36–51.

Duan, Y., Zhang, Y., Dong, H., Wang, Y. and Zhang, J., 2017. Effect of the dietary probiotic Clostridium butyricum on growth, intestine antioxidant capacity and resistance to high temperature stress in kuruma shrimp Marsupenaeus japonicus. Journal of Thermal Biology, 66, pp. 93–100.

Fallet, M., Montagnani, C., Petton, B., Dantan, L., De Lorgeril, J., Comarmond, S., Chaparro, C., Toulza, E., Boitard, S., Escoubas, J.M. and Vergnes, A., 2022. Early life microbial exposures shape the *Crassostrea gigas* immune system for lifelong and intergenerational disease protection. Microbiome, 10(1), pp. 85.

Froelich, B., Ayrapetyan, M. and Oliver, J.D., 2013. Integration of *Vibrio vulnificus* into marine aggregates and its subsequent uptake by *Crassostrea virginica* oysters. Applied and environmental microbiology, 79(5), pp.1454–1458.

Garnier, M., Labreuche, Y., Garcia, C., Robert, M. and Nicolas, J.L., 2007. Evidence for the involvement of pathogenic bacteria in summer mortalities of the Pacific oyster *Crassostrea gigas*. Microbial ecology, 53, pp. 187–196.

Garnier, M., Labreuche, Y. and Nicolas, J.L., 2008. Molecular and phenotypic characterization of *Vibrio aestuarianus* subsp. *francensis* subsp. nov., a pathogen of the oyster *Crassostrea gigas*. Systematic and applied microbiology, 31(5), pp. 358–365.

Geng, H. and Belas, R., 2010. Expression of tropodithietic acid biosynthesis is controlled by a novel autoinducer. Journal of Bacteriology, 192(17), pp. 4377–4387.

Geng, H., Bruhn, J.B., Nielsen, K.F., Gram, L. and Belas, R., 2008. Genetic dissection of tropodithietic acid biosynthesis by marine roseobacters. Applied and environmental microbiology, 74(5), pp. 1535–1545.

Gensollen, T., Iyer, S.S., Kasper, D.L. and Blumberg, R.S., 2016. How colonization by microbiota in early life shapes the immune system. Science, 352(6285), pp. 539–544.

Goudenège, D., Travers, M.A., Lemire, A., Petton, B., Haffner, P., Labreuche, Y., Tourbiez, D., Mangenot, S., Calteau, A., Mazel, D. and Nicolas, J.L., 2015. A single regulatory gene is sufficient to alter *Vibrio aestuarianus* pathogenicity in oysters. Environmental Microbiology, 17(11), pp. 4189–4199.

Green, T.J., Siboni, N., King, W.L., Labbate, M., Seymour, J.R. and Raftos, D., 2019. Simulated marine heat wave alters abundance and structure of *Vibrio* populations associated with the Pacific Oyster resulting in a mass mortality event. Microbial ecology, 77, pp. 736–747.

Hartig, F., 2024. DHARMa: Residual Diagnostics for Hierarchical (Multi-Level / Mixed) Regression Models. R package version 0.4.7. Available at: https://CRAN.R-project.org/package=DHARMa.

Harvell, C.D., Kim, K., Burkholder, J.M., Colwell, R.R., Epstein, P.R., Grimes, D.J., Hofmann, E.E., Lipp, E.K., Osterhaus, A.D.M.E., Overstreet, R.M. and Porter, J.W., 1999. Emerging marine diseases--climate links and anthropogenic factors. Science, 285(5433), pp. 1505–1510.

Helm, M.M., Bourne, N. and Lovatelli, A., 2004. The hatchery culture of bivalves: a practical manual. FAO.

Henriksen, N.N.S.E., Schostag, M.D., Balder, S.R., Bech, P.K., Strube, M.L., Sonnenschein, E.C. and Gram, L., 2022. The ability of *Phaeobacter inhibens* to produce tropodithietic acid influences the community dynamics of a microalgal microbiome. ISME Communications, 2(1), pp. 109.

Huang, B., Zhang, L., Li, L., Tang, X. and Zhang, G., 2015. Highly diverse fibrinogen-related proteins in the Pacific oyster *Crassostrea gigas*. Fish & Shellfish Immunology, 43(2), pp. 485–490.

Joo, H.S., Fu, C.I. and Otto, M., 2016. Bacterial strategies of resistance to antimicrobial peptides. Philosophical Transactions of the Royal Society B: Biological Sciences, 371(1695).

Kach, D.J. and Ward, J.E., 2008. The role of marine aggregates in the ingestion of picoplankton-size particles by suspension-feeding molluscs. Marine Biology, 153(5), pp.797–805.

Karim, M., Zhao, W., Rowley, D., Nelson, D. and Gomez-Chiarri, M., 2013. Probiotic strains for shellfish aquaculture: protection of eastern oyster, *Crassostrea virginica*, larvae and juveniles against bacterial challenge. Journal of Shellfish Research, 32(2), pp. 401–408.

Kassambara, A., Kosinski, M. and Biecek, P., 2024. survminer: Drawing Survival Curves using ‘ggplot2’. R package version 0.5.0. Available at: https://CRAN.R-project.org/package=survminer.

Katoh, K., Misawa, K., Kuma, K., & Miyata, T., 2002. MAFFT: A novel method for rapid multiple sequence alignment based on fast Fourier transform. Nucleic Acids Research, 30(14), pp. 3059–3066. 10.1093/nar/gkf436

Kesarcodi-Watson, A., Miner, P., Nicolas, J.L. and Robert, R., 2012. Protective effect of four potential probiotics against pathogen-challenge of the larvae of three bivalves: Pacific oyster (*Crassostrea gigas*), flat oyster (*Ostrea edulis*) and scallop (*Pecten maximus*). Aquaculture, 344, pp. 29–34.

King, W.L., Siboni, N., Williams, N.L., Kahlke, T., Nguyen, K.V., Jenkins, C., Dove, M., O’Connor, W., Seymour, J.R. and Labbate, M., 2019. Variability in the composition of Pacific oyster microbiomes across oyster families exhibiting different levels of susceptibility to OsHV-1 μvar disease. Frontiers in microbiology, 10, p.473.

Kohl, K.D., 2025. Through the microbial looking glass: our shifting understanding of the holobiont and microbes as mediators of organismal biology. Integrative and Comparative Biology, 65(3), pp.727–735.

Kozlov, A. M., Darriba, D., Flouri, T., Morel, B., & Stamatakis, A., 2019. RAxML-NG: A fast, scalable and user-friendly tool for maximum likelihood phylogenetic inference. Bioinformatics, 35(21), pp. 4453–4455. 10.1093/bioinformatics/btz305

Lannig, G., Flores, J.F. and Sokolova, I.M., 2006. Temperature-dependent stress response in oysters, *Crassostrea virginica*: pollution reduces temperature tolerance in oysters. Aquatic toxicology, 79(3), pp. 278–287.

Lee, S.Y., Park, S., Oh, T.K. and Yoon, J.H., 2012. *Celeribacter baekdonensis* sp. nov., isolated from seawater, and emended description of the genus *Celeribacter* Ivanova et al. 2010. International journal of systematic and evolutionary microbiology, 62(Pt_6), pp. 1359–1364.

Lenth, R., 2024. emmeans: Estimated Marginal Means, aka Least-Squares Means. R package version 1.10.6. Available at: https://CRAN.R-project.org/package=emmeans.

Lehtinen, R.M., 2023. Empirical evidence for the rescue effect from a natural microcosm. Animals, 13(12), p.1907.

Letunic, I. and Bork, P., 2024. Interactive Tree of Life (iTOL) v6: Recent updates to the phylogenetic tree display and annotation tool. Nucleic Acids Research, 52(W1), W78–W82. 10.1093/nar/gkae268

Li, Y.F., Xu, J.K., Chen, Y.W., Ding, W.Y., Shao, A.Q., Liang, X., Zhu, Y.T. and Yang, J.L., 2019. Characterization of gut microbiome in the mussel *Mytilus galloprovincialis* in response to thermal stress. Frontiers in physiology, 10, pp. 1086.

Li, J., Zou, Y., Li, Q., Zhang, J., Bourne, D.G., Lyu, Y., Liu, C. and Zhang, S., 2023. A coral-associated actinobacterium mitigates coral bleaching under heat stress. Environmental Microbiome, 18(1), pp. 83.

Liang, L., 2003. Investigation of secondary metabolites of North Sea bacteria: fermentation, isolation, structure elucidation and bioactivity. Georg-August-Universitaet Goettingen (Germany).

Lokmer, A. and Wegner, K.M., 2015. Hemolymph microbiome of Pacific oysters in response to temperature, temperature stress and infection. The ISME journal, 9(3), pp. 670–682.

Luo, H. and Moran, M.A., 2014. Evolutionary ecology of the marine Roseobacter clade. Microbiology and Molecular Biology Reviews, 78(4), pp. 573–587.

Lupo, C., Travers, M.A., Tourbiez, D., Barthélémy, C.F., Beaunée, G. and Ezanno, P., 2019. Modeling the transmission of *Vibrio aestuarianus* in pacific oysters using experimental infection data. Frontiers in Veterinary Science, 6, pp. 142.

Mackenzie, C.L., Pearce, C.M., Leduc, S., Roth, D., Kellogg, C.T., Clemente-Carvalho, R.B. and Green, T.J., 2022. Impacts of seawater pH buffering on the larval microbiome and carry-over effects on later-life disease susceptibility in Pacific oysters. Applied and Environmental Microbiology, 88(22), pp.e01654–22.

Malham, S.K., Cotter, E., O’Keeffe, S., Lynch, S., Culloty, S.C., King, J.W., Latchford, J.W. and Beaumont, A.R., 2009. Summer mortality of the Pacific oyster, Crassostrea gigas, in the Irish Sea: the influence of temperature and nutrients on health and survival. Aquaculture, 287(1-2), pp.128–138.

Margulis, L., 1993. Symbiosis in cell evolution: Microbial communities in the archean and proterozoic eons. New York, NY: WH Freeman and Co.

McFall-Ngai, M., Hadfield, M.G., Bosch, T.C., Carey, H.V., Domazet-Lošo, T., Douglas, A.E., Dubilier, N., Eberl, G., Fukami, T., Gilbert, S.F. and Hentschel, U., 2013. Animals in a bacterial world, a new imperative for the life sciences. Proceedings of the National Academy of Sciences, 110(9), pp. 3229–3236.

McMurdie, P.J. and Holmes S., 2013. phyloseq: An R package for reproducible interactive analysis and graphics of microbiome census data. PLoS ONE, 8(4).

Mou, X., Moran, M.A., Stepanauskas, R., González, J.M. and Hodson, R.E., 2005. Flow-cytometric cell sorting and subsequent molecular analyses for culture-independent identification of bacterioplankton involved in dimethylsulfoniopropionate transformations. Applied and environmental microbiology, 71(3), pp. 1405–1416.

Moradi, M., Magalhaes, P.R., Peixoto, R.S., Jonck, C.C., François, D., Bellot, A.C.F., Teixeira, J.B., Silveira, C.S., Duarte, G., Evangelista, H. and Barbosa, C.F., 2023. Roseobacter-enriched mitigate thermal stress-and pathogen-driven impacts on coral skeleton. Frontiers in Marine Science, 10, pp. 1212690.

Nyholm, S.V. and Graf, J., 2012. Knowing your friends: invertebrate innate immunity fosters beneficial bacterial symbioses. Nature Reviews Microbiology, 10(12), pp. 815–827.

Oksanen, J., Blanchet, F.G., Kindt, R., Legendre, P., Minchin, P.R., O’hara, R.B., Simpson, G.L., Solymos, P., Stevens, M.H.H., Wagner, H. and Oksanen, M.J., 2013. Package ‘vegan’. Community ecology package, version, 2(9), pp.1–295.

Paradis E. and Schliep K., 2019. ape 5.0: an environment for modern phylogenetics and evolutionary analyses in R.” Bioinformatics, 35, pp. 526–528.

Pierce, M.L. and Ward, J.E., 2019. Gut microbiomes of the eastern oyster (*Crassostrea virginica*) and the blue mussel (*Mytilus edulis*): temporal variation and the influence of marine aggregate-associated microbial communities. Msphere, 4(6), pp.10–1128.

Pujalte, M.J., Lucena, T., Ruvira, M.A., Arahal, D.R. and Macián, M.C., 2014. 20 The Family Rhodobacteraceae.

R Core Team, 2025. R: A Language and Environment for Statistical Computing. R Foundation for Statistical Computing, Vienna, Austria. https://www.R-project.org/

Ranson, H.J., LaPorte, J., Spinard, E., Gomez-Chiarri, M., Nelson, D.R. and Rowley, D.C., 2018. Draft Genome Sequence of *Loktanella maritima* Strain YPC211, a Commensal Bacterium of the American Lobster (*Homarus americanus*). Genome Announcements, 6(18), pp. 10–1128.

Ranson, H.J.G., 2020. Chemical Investigation of Bacterial Interactions Involving Pathogens (Doctoral dissertation, University of Rhode Island).

Richards, G.P., Watson, M.A., Needleman, D.S., Church, K.M. and Häse, C.C., 2015. Mortalities of Eastern and Pacific oyster larvae caused by the pathogens *Vibrio coralliilyticus* and *Vibrio tubiashii*. Applied and Environmental Microbiology, 81(1), pp.292–297.

Risely, A., 2020. Applying the core microbiome to understand host–microbe systems. Journal of Animal Ecology, 89(7), pp. 1549–1558.

Rook, G., Martinelli, R. and Brunet, L., 2005. The “Old Friends” hypothesis: How early contact with certain microorganisms may influence immunoregulatory circuits. Perinatal programming. Boca Raton: CRC Press, pp. 183–194.

Rosado, P.M., Leite, D.C., Duarte, G.A., Chaloub, R.M., Jospin, G., Nunes da Rocha, U., Saraiva, J.P., Dini-Andreote, F., Eisen, J.A., Bourne, D.G. and Peixoto, R.S., 2019. Marine probiotics: increasing coral resistance to bleaching through microbiome manipulation. The ISME journal, 13(4), pp. 921–936.

Rosani U, Bai CM, Maso L, Shapiro M, Abbadi M, Domeneghetti S, Wang CM, Cendron L, MacCarthy T, and Venier P., 2019. A-to-I editing of *Malacoherpesviridae* RNAs supports the antiviral role of ADAR1 in mollusks. BMC Evolutionary Biology, 19(1), pp. 149.

Rosenthal, J.J. and Eisenberg, E., 2023. Extensive recoding of the neural proteome in cephalopods by RNA editing. Annual Review of Animal Biosciences, 11, pp. 57–75.

Ruiz-Ponte, C., Samain, J.F., Sanchez, J.L. and Nicolas, J.L., 1999. The benefit of a Roseobacter species on the survival of scallop larvae. Marine Biotechnology, 1(1), pp. 52–59.

Scanes, E., Siboni, N., Rees, B. and Seymour, J.R., 2023. Acclimation in intertidal animals reduces potential pathogen load and increases survival following a heatwave. Iscience, 26(6).

Schliep, K., Potts, A. J., Morrison, D. A., Grimm, G. W., 2017, Intertwining phylogenetic trees and networks. Methods in Ecology and Evolution, 8, pp. 1212–1220. doi: 10.1111/2041-210X.12760

Seemann, T. (2014). Prokka: Rapid prokaryotic genome annotation. *Bioinformatics (Oxford*, England), 30(14), pp. 2068–2069. 10.1093/bioinformatics/btu153

Sharifah, E.N. and Eguchi, M., 2011. The phytoplankton *Nannochloropsis oculata* enhances the ability of Roseobacter clade bacteria to inhibit the growth of fish pathogen *Vibrio anguillarum*. PLoS One, 6(10), pp.e26756.

Song, X., Wang, H., Xin, L., Xu, J., Jia, Z., Wang, L. and Song, L., 2016. The immunological capacity in the larvae of Pacific oyster *Crassostrea gigas*. Fish & shellfish immunology, 49, pp. 461–469.

Sorée, M., Delavat, F., Lambert, C., Lozach, S., Papin, M., Petton, B., Passerini, D., Dégremont, L. and Hervio Heath, D., 2022. Life history of oysters influences *Vibrio parahaemolyticus* accumulation in Pacific oysters (*Crassostrea gigas*). Environmental Microbiology, 24(9), pp. 4401–4410.

Surry, L.B., Sutherland, B.J., Lunda, S.L., Loudon, A.H., Divilov, K., Langdon, C.J. and Green, T.J., 2024. The presence of the Pacific Oyster OSHV-1 resistance marker on chromosome 8 does not impact susceptibility to infection by *Vibrio aestuarianus*. bioRxiv.

Therneau TM, 2024. coxme: Mixed Effects Cox Models. R package version 2.2-22, https://CRAN.R-project.org/package=coxme.

Unzueta-Martínez, A., Welch, H. and Bowen, J.L., 2022. Determining the composition of resident and transient members of the oyster microbiome. Frontiers in Microbiology, 12, pp. 828692.

Unzueta-Martínez, A., Scanes, E., Parker, L.M., Ross, P.M., O’Connor, W. and Bowen, J.L., 2022. Microbiomes of the Sydney Rock Oyster are acquired through both vertical and horizontal transmission. Animal Microbiome, 4(1), pp. 32.

Wick, R. R., Judd, L. M., Gorrie, C. L., & Holt, K. E. (2017). Unicycler: Resolving bacterial genome assemblies from short and long sequencing reads. PLOS Computational Biology, 13(6), e1005595. 10.1371/journal.pcbi.1005595

Wilson, M.Z., Wang, R., Gitai, Z. and Seyedsayamdost, M.R., 2016. Mode of action and resistance studies unveil new roles for tropodithietic acid as an anticancer agent and the γ-glutamyl cycle as a proton sink. Proceedings of the National Academy of Sciences, 113(6), pp.1630–1635.

Wirth, J.S. and Whitman, W.B., 2018. Phylogenomic analyses of a clade within the roseobacter group suggest taxonomic reassignments of species of the genera *Aestuariivita*, *Citreicella*, *Loktanella*, *Nautella, Pelagibaca, Ruegeria, Thalassobius, Thiobacimonas* and *Tropicibacter*, and the proposal of six novel genera. International journal of systematic and evolutionary microbiology, 68(7), pp. 2393–2411.

Wood, S.N., 2011. Fast stable restricted maximum likelihood and marginal likelihood estimation of semiparametric generalized linear models. Journal of the Royal Statistical Society (B), 73(1), pp.3–36

Wright ES, 2016. Using DECIPHER v2.0 to Analyze Big Biological Sequence Data in R. The R Journal, 8(1), pp. 352–359.

Yilmaz, P., Parfrey, L.W., Yarza, P., Gerken, J., Pruesse, E., Quast, C., Schweer, T., Peplies, J., Ludwig, W. and Glöckner, F.O., 2014. The SILVA and “all-species living tree project (LTP)” taxonomic frameworks. Nucleic acids research, 42(D1), pp. 643–648.

